# Birdwatchers’ resilience to perturbation in India buffers citizen science from pandemic-induced biases

**DOI:** 10.1101/2023.12.11.571083

**Authors:** Karthik Thrikkadeeri, Ashwin Viswanathan

**Affiliations:** Nature Conservation Foundation

**Keywords:** Birdwatcher behaviour, COVID-19 pandemic, data characteristics, eBird, effort

## Abstract

Most systematic projects to monitor bird populations, like breeding bird surveys, require large and coordinated volunteer networks that are lacking in many parts of the world such as the Global South. Data from less systematic citizen science (CS) programmes offer an alternative to data from systematic initiatives in these regions, but the semi-structured nature of such data also presents several challenges. The utility of semi-structured CS data to monitor bird species abundance is contingent on how, where, and how comparably birdwatchers watch birds, year on year. Trends inferred directly from the data can be confounded during years when birdwatchers may behave differently, such as during the COVID-19 pandemic. We wanted to ascertain how the data uploaded from India to one such CS platform, eBird, was impacted by this deadliest global pandemic of the 21st century. To understand whether eBird data from the pandemic years in India is useful and comparable to data from adjacent years, we explored several quantitative and qualitative aspects of the data (such as birdwatcher behaviour) at multiple spatial and temporal scales. We found no negative impact of the pandemic on data generation. Data characteristics changed largely only during the peak pandemic months characterised by high fatality rates and strict lockdowns, possibly due to decreased human mobility and social interaction. It remained similar to the adjacent years during the rest of this restrictive period, thereby reducing the impact of the aberrant peak months on any annual inference. Moreover, impacts on data characteristics varied widely across states in India, resulting in no strong consistent trend at the national level—unlike results from elsewhere in the world. Our findings show that birdwatchers in India as contributors to CS were resilient to disturbance, and that the effects of the pandemic on birdwatching effort and birdwatcher behaviour are highly scale- and context-dependent. In summary, eBird data in India from the pandemic years remains useful and interpretable for most large-scale applications, such as abundance trend estimation, but will benefit from preliminary data quality checks when utilised at a fine scale.

**Lay summary:**

- Citizen science platforms like eBird comprise vast repositories of data generated by casual birdwatching.
- Such data are vital to understanding bird population trends, but their usability reduces when birdwatchers change where, when and how much they watch birds from year to year.
- Given the impact of the COVID-19 pandemic on our everyday lives, we wondered whether it also impacted the way people reported birds, thereby reducing the usability of the data in trend analyses.
- We analysed data uploaded to eBird from India, and found that the impacts of the pandemic on this data were largely restricted to April and May in 2020, and to a lesser
- extent in 2021. During these months that coincided with the greatest health impacts, birdwatchers avoided travel, groups and public spaces.
- Birdwatchers were resilient; they bounced back soon after these difficult periods, and started birding like they had done before the pandemic.
- Because the impact was limited to short periods and few regions, we conclude that eBird data from India during the pandemic still remains useful for analyses of bird abundance trends.

## 1 Introduction

### 1.1 Birds and citizen science

Citizen science (CS) today is a research approach that engages volunteers from the general public—most of whom are not trained as scientists—in collecting and/or analysing and interpreting data as a part of scientific enquiry (<u>Silvertown 2009</u>). CS programmes are particularly effective when they revolve around popular hobbies or activities that people enjoy, such as birdwatching. Birdwatching (hereafter birding) is a nature-based activity that is enjoyed by a large number of people around the world, making CS projects involving birds especially appealing to the public. Birders, as a matter of practice, make “checklists” of the bird species they observe during birding sessions. By pooling this data into a common public database, numerous birders with varying degrees of experience—from trained researchers to seasoned birders to laypersons—are able to contribute to the scientific study of birds.

Systematic CS bird monitoring programmes (see Miller-Rushing et al. 2012) that engage birders are resource- and volunteer-intensive (e.g., Greenwood et al. 1995, Sauer et al. 2013, BirdLife Australia 2015, Praveen et al. 2022, Swiss Ornithological Institute 2023). In parts of the world like India where this is not yet feasible at large scales, CS projects with “semi-structured” protocols (Pocock et al. 2017, Di Cecco et al. 2021) work better. For instance, eBird (Sullivan et al. 2014) is widely used by birders in India and enables large-scale robust scientific analysis and output (e.g., SoIB 2023). eBird data has proven very useful in better understanding bird ecology, species ranges (e.g., Sreekumar and Nameer 2021, Freeman et al. 2022), migration (e.g., Youngflesh et al. 2021, Menon et al. 2023), behaviour (e.g., Winger et al. 2019, Alaniz et al. 2020), morphology (e.g., Vrettos et al. 2021, Menon et al. 2022) and conservation action (e.g., Sullivan et al. 2017, Ruiz-Gutierrez et al. 2021, SoIB 2023).

However, a drawback is that eBird checklists vary in aspects such as effort and the data is not homogeneous in space and time (as there is no single fixed protocol), thereby creating noticeable biases (<u>Johnston et al. 2021</u>). Largely associated with birder behaviour, such biases make the data difficult to use to inform yearly abundance trends. Although some recent analyses show that appropriate statistical adjustments can reduce these biases such that results closely match those from systematic data (<u>Johnston et al. 2021</u>, <u>Viswanathan n.d.</u>, <u>Viswanathan et al. n.d.</u>), biases in semi-structured data become especially pronounced when there are large-scale changes in human (birder) behaviour, such as presumably during the COVID-19 pandemic (see next). Given the widespread utility of semi-structured CS data today for conservation, an urgent question is whether the pandemic has rendered eBird data from the pandemic years unusable in combination with the rest of the dataset for estimating yearly abundance trends.

### 1.2 Birding in the time of COVID-19

COVID-19 is a highly infectious disease that resulted in the deadliest pandemic of the 21st century, with a death toll of more than 6.9 million (around 0.1% of the global population; Wikipedia 2023; Worldometer <u>2023</u>). Governments across the world therefore took various precautionary and preventive measures, and implemented restrictions such as nation- or state-wide lockdowns and curfews, quarantines and self-isolation, and shutdown of non-essential public services and activities (see <u>Hale et al. 2020</u>). Given that the eBird enterprise hinges entirely on public participation, this widespread public upheaval might have changed the very nature of eBird data during this period.

#### 1.2.1 How much birding?

The COVID-19 pandemic has generally posed a big challenge for CS programmes (e.g., <u>Hochachka et al. 2021</u>, <u>Kishimoto and Kobori 2021</u>). Social and travel restrictions, in combination with physical and mental distress (<u>Li et al. 2020</u>, <u>Van Bavel et al. 2020</u>), may have led to a decline in the amount of hobbyist as well as systematic birding, and therefore in the quantity of data generated. Some CS projects did indeed show evidence of reduced participation and data generation (e.g., <u>Kishimoto and Kobori 2021</u>), but others in fact witnessed an increase in data generated (e.g., <u>Basile et al. 2021</u>). Outdoor recreational activities such as birding are known to restore attention and play a vital role in improving physical and mental health (<u>Weng and Chiang 2014</u>). The pandemic brought an increased desire in people to connect with the natural world (<u>Marsh et al. 2021</u>), as well as active promotion and encouragement of nature-related activities such as birding (e.g., <u>Bird Count India 2020</u>). Thus, many turned to birding as their activity of choice (<u>Randler et al. 2020</u>).

#### 1.2.2 What kinds of birding?

Regardless of how much more or less data was generated during the pandemic, the characteristics of this data could also have changed. For example, Randler et al. (<u>2020</u>) found differences in aspects of birder behaviour, such as who was birding and with whom, as well as where and when birding (and thus data generation) was happening. Restricted mobility has led to data being less spread out in space, with observations being concentrated in urbanised locations (<u>Vardi et al. 2021</u>) and covering a lower overall diversity of habitats (<u>Hochachka et al. 2021</u>), thus reducing its informational value (<u>Zhang and Zhu 2020</u>). On the other hand, Kishimoto and Kobori (<u>2021</u>) found that the restrictions resulted in lower clustering of data, potentially because birders were limited to their erstwhile under-surveyed neighbourhoods and could not upload as many observations as usual from birding hotspots (either urban or rural).

#### 1.2.3 Scale and generalisability

Hochachka et al. (<u>2021</u>) studied changes in birder behaviour during a month-long lockdown across three countries: USA (states of California and New York), Spain and Portugal. The changes were not as substantial as expected from the magnitude of change to daily life. Notably, they found differential responses to the pandemic in the respective birding communities, and suggested that such differences are likely to exist amongst geographical units across the world, thus cautioning against a global generalisation of how the pandemic influenced birding and birder behaviour. Moreover, such studies of the pandemic impacts have not explored broader temporal scales—time scales relevant for large-scale analyses of abundance trends, such as those contained in the State of India’s Birds report (<u>SoIB 2023</u>).

### 1.3 Our aim

India’s rapid recent progress in volunteer-driven bird monitoring is both unique and pioneering, in the context of the Global South where large-scale resource-intensive systematic surveys may not be possible in the near future. The State of India’s Birds reports (<u>SoIB 2020</u>, <u>2023</u>) have demonstrated that semi-structured monitoring can be immensely valuable for assessing the health of a region’s birds. Because such assessments are intended to inform conservation policy, there is an urgent need to ensure robustness of the data from birding to disruptive events like the COVID-19 pandemic or even potential future climate crises.

Birders in India and elsewhere in the Global South, under such circumstances, might behave differently from those in North America and Europe, and therefore respond differently to events such as the COVID-19 pandemic, due to different socioeconomic realities, aspirations, and politics. For instance, adherence to social distancing measures rapidly declined across socioeconomic strata in India after the initial months of lockdowns (<u>Schaner et al. 2022</u>). The epidemiological and welfare value of such measures in countries such as India is often lower than in higher-income countries due to the economic sacrifices involved (<u>Barnett-Howell et al. 2021</u>).

In this study, we sought a comprehensive understanding of how birders in India may have changed their behaviour during the pandemic, and what this may indicate about the utility of the generated data. We specifically focused on the use of the platform eBird in India, and asked whether or not eBird data characteristics were similar before, during and after the pandemic. Where dissimilar, we then tried to ascertain the extent of these changes, by assessing transience (peak pandemic months vs years) and spatial variation (states vs country). Finally, we use this understanding to synthesise the implications of the pandemic on the utility of eBird data for long-term bird monitoring—specifically for abundance trend estimation in India—and also outline guidelines for using this data for such applications.

## 2 Methods

### 2.1 Data preparation

We used the “relMay-2022” version of the publicly available <u>eBird Basic Dataset (EBD)</u> for India, containing all data uploaded until 31 May 2022 (downloaded on 15 June 2022, follows taxonomy in <u>Clements et al. 2019</u>). We combined <u>sensitive species data</u>, which is not included in the public download, after requesting this separately from eBird. Before analysis, we applied a set of filters to standardise the dataset (Supplementary Material Section S1.1.1), so as to prevent any extreme and outlying cases from skewing the main patterns. Our final eBird dataset contained 814,524 unique “qualifying checklists”, i.e., lists that passed the filters.

All analyses presented in this study use “migratory year” (MY; June of one calendar year to May of the next; see Supplementary Material Section S1.1.1 for rationale) as a unit time period and we analysed data over a period of four migratory years, from 1 June 2018 to 31 May 2022 (Fig. 1). We consider a relatively fine temporal scale of months in order to account for within-year variations in bird and birder behaviour. Moreover, in India severe COVID-19 restrictions and lockdowns were limited to a few months in 2020 and 2021 (<u>The Economic Times 2020</u>, <u>The Indian Express 2021</u>), and this scale allows us to consider these patterns separately from patterns in other months of the “pandemic years” (the migratory years 2019– 20 and 2020–21, coded as “DUR”; 2018–19 and 2021–22 coded as “BEF” and “AFT” respectively).

**Figure 1:**
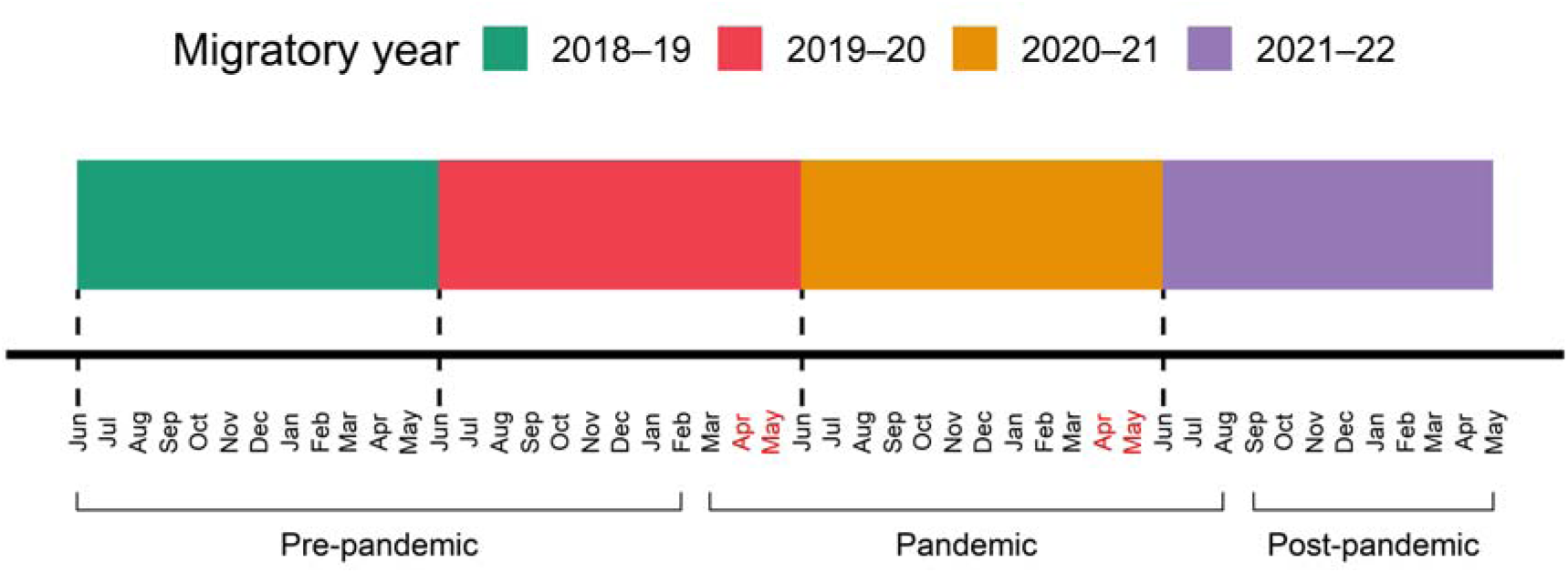
Timeline of the study period. The spans of the four migratory years are shown in different colours, and pre-pandemic, pandemic and post-pandemic months are also signified. April and May of both pandemic years (shown in red text) are considered as the peak pandemic months.

We gridded the spatial extent of the country into 0.225° X 0.225° cells (20–24.8 km X 25 km, depending on latitude) and used the latitude-longitude information associated with each of the qualifying checklists to place them in corresponding grid cells. We also classified the checklists as “urban” or “non-urban” using remote sensing data on land cover (MODIS Land Cover Type Product MCD12Q1, <u>Friedl and Sulla-Menashe 2019</u>) with the help of the *luna* (<u>Ghosh et al. 2020</u>) package (see Supplementary Material Section S1.1.2 for details). The spatial resolution of our grid is large (relative to practices elsewhere like North America), and we avoided associating eBird checklist locations with remotely sensed habitat variables, because of the significant inherent imprecision in where and how checklist locations are assigned (e.g., eBird hotspots often encompass large areas and numerous habitats).

### 2.2 Analysis of data characteristics

To analyse changes in the nature of eBird data, we devised a suite of metrics (see below) that help gauge various characteristics of the data. These metrics were all calculated per month, across the four years, and these values were compared to investigate changes in data characteristics over time. Except for data quantity (Section 2.2.1) and spatial spread (Section 2.2.5), all the metrics were analysed at two spatial scales: for the whole country and for each of four states. The four states we considered in our study in order to highlight nuances were Karnataka, Kerala, Maharashtra and Assam. These states have relatively high eBirding activity and cover different regions of the country, but also differ in characteristics of the birding community as well as in response to the pandemic owing to differences in local governance, policy and general public perspective.

The metrics of birder behaviour (Section 2.2.2), urban bias (its additive inverse; Section 2.2.3) and spatial coverage (Section 2.2.4) have a common directionality: if the pandemic did indeed change these characteristics of the data, we expected the values of each of these metrics to decline from pre-pandemic levels. The metrics are described below, and the process of summarising them and interpreting their combined meaning is detailed in Section 2.2.6.

#### 2.2.1 Data quantity

To test whether the pandemic led to an increase or decrease in eBird data quantity, we explored two separate aspects of quantity: data volume and participation. We compared the number of qualifying lists and the number of active eBirders respectively for every month across our years of interest, using simple summaries and visualisations.

#### 2.2.2 Metrics of birder behaviour

1. *Group birding*: Were there fewer instances of group birding during the pandemic? For this, we calculated the proportion of an observer’s total uploaded qualifying lists that were instances of group birding, defined as lists with three or more observers as specified in the “party size” field.
2. *Site fidelity*: Given the various restrictions on movement and travel, did birders selectively visit certain areas more often than they previously did, and therefore visit fewer areas overall? In other words, did birders show higher “site fidelity” during the pandemic? We calculated the number of grid cells visited by each birder, i.e., the inverse of site fidelity.
3. *Hotspot birding*: eBird “hotspots” are public birding locations demarcated by eBird users to aggregate and display data from multiple birders (<u>eBird 2022</u>), and are different from “personal” birding locations which are usually visited by just one or few birders. Given the overall decline in public/social interactions during the pandemic, did people also tend to avoid birding in eBird hotspots? We calculated, out of an observer’s total uploaded qualifying lists, the proportion that was from eBird hotspots.
4. *Birding protocol*: “Traveling” and “Stationary” are the two most commonly followed protocols in eBird observations. Considering the restrictions on movement during the pandemic, we asked whether birders made fewer travelling lists (which we defined as lists using the “Traveling” protocol and covering > 0.3 km), instead preferring stationary lists (from, say, their homes). We calculated, out of an observer’s total uploaded qualifying lists, the proportion that was of travelling lists.
5. *List distance*: Among the observations that followed the “Traveling” protocol, we asked whether travelling distances were overall shorter during the pandemic. Here, we simply used the distance travelled, measured as part of the checklist-level effort metadata.
6. *List duration*: Did eBirders make shorter-duration checklists during the pandemic? Here, we considered checklist-level effort using checklist duration from the effort metadata.
7. *List length*: List length is defined as the total number of species observed in a checklist. This has been used as a metric of effort in some studies (e.g., <u>Szabo et al. 2010</u>, <u>Isaac et al. 2014</u>), and is often used as a substitute for list duration or distance in such analyses (e.g., <u>SoIB 2023</u>). We used list length to answer, from a different angle, the question of whether checklists became shorter during the pandemic.

#### 2.2.3 Urban bias

A majority of eBird users are likely to be urban-dwelling, both in India and across the world, but often move far and wide for birding. Birders from urban areas may therefore account for a large proportion of the birding in non-urban areas as well. With the pandemic limiting movement, did urban areas see an increase in birding while non-urban areas saw the opposite trend? If so, this would have implications for how well these two areas are represented in the data compared to other years, potentially biasing analyses. We calculated the proportion of lists out of an observer’s total uploaded qualifying lists that were from “urban” sites (as defined previously) to estimate urban bias. For the composite analysis (Section 2.2.6), we used the additive inverse of this metric, i.e., non-urban bias.

#### 2.2.4 Spatial coverage

Given the restrictions on travel, a decrease in overall spatial coverage of the data from the pandemic was a major concern, because the country as a whole would be less well represented than in other years, thus biasing comparisons. We calculated the proportion of grid cells across the country that were covered in each time period. The information threshold for a grid cell to be considered “covered” was set at 5 qualifying checklists (per month per year).

#### 2.2.5 Spatial spread

Did the pandemic create spatial clusters of positive or negative change in birding activity across grid cells in the country, or were changes in birding (if any) relatively uniformly spread out? If strong changes (positive or negative) were concentrated in specific parts of the country, data from the pandemic would be biased because they do not represent well those regions in the country that are typically sampled by birders. For each grid cell, we calculated proportional change in number of qualifying lists uploaded for transition periods (BEF–DUR and DUR–AFT, where DUR was an aggregate of MY 2019–20 and 2020–21). Here, the larger scale was our temporal unit of focus (i.e., years, not months), and the analysis was only done for the entire country. See Supplementary Material Section S1.2 for details on how this metric was calculated and interpreted.

#### 2.2.6 Analysis

All metrics except data quantity and spatial spread were analysed using the formal mixed-effects model framework, mostly using Generalised Linear Mixed Models (GLMMs, see <u>Bolker et al. 2009</u>). We used the “glmer()” function in the *lme4* package (<u>Bates et al. 2015</u>) for the R environment (<u>R Core Team 2021</u>) to fit GLMMs. Observer identity defined by unique observer number was used as the random effect for the metrics of birder behaviour and urban bias; spatial identity defined by grid cell ID was used as the random effect for spatial coverage (see Supplementary Material Section S1.3). All GLMMs were fitted with binomial error distribution and cloglog link function, using BOBYQA optimiser.

Some metrics like site fidelity, list distance, list duration and list length were either whole-number or continuous numerical variables that did not include zero in the range of possible values. For these, since GLMMs with binomial or Poisson error distributions are not appropriate, we instead fitted LMMs (Linear Mixed Models, “lmer()” function in *lme4*) with the same fixed and random effects specifications and using BOBYQA optimiser, after first log-transforming (natural log) the response variables to approximate normality. The specifications of the various models of data metrics are listed in Supplementary Material Section S1.3.

For all models, model diagnostics were inspected using the *DHARMa* package (<u>Hartig 2022</u>), to ensure that there were no major violations of model assumptions. We used the “bootMer()” function in *lme4* in order to test the significance of the fitted model results, by obtaining mean and CI estimates using semi-parametric bootstrapping. The bootstrapping was done with 1000 simulations and with parallel core processing (with the help of packages *parallel*, <u>R Core Team 2021</u>, and *VGAM*, <u>Yee 2010</u>).

The model results of the nine such metrics of data characteristics were summarised as follows to give an understanding of the overall impact of the pandemic (and its scale) on birding and birder behaviour in India. Model predictions for each metric (per month and year) were first transformed into proportional change relative to corresponding months in MY 2018–19, while also transforming the associated uncertainty appropriately. In the case of urban bias, before transforming to proportional change, we initially calculated the additive inverse of predicted values (i.e., non-urban bias), in order to match our expectation of decline during the pandemic. After the transformation, the months of a year were classified into whether they were peak impact months (April–May) or not, and values within these were averaged across individual months and metrics, accompanied by propagation of uncertainty. Thus, we finally obtained measures of proportional change in data quality metrics (and associated confidence intervals) for each year and month category, relative to the standard of MY 2018–19, which allowed us to infer both magnitude and scale of the pandemic impact. Due to the common expected directionality in each of the nine metrics, we expected the pandemic impact to manifest as a decline from pre-pandemic levels when the metrics were combined into this single index. The absence of any perceivable difference in a given period would indicate that there is no evidence of fundamental characteristics of the data having changed during that period.

### 2.3 Software

All analyses were performed in the R environment (<u>R Core Team 2021</u>) in RStudio (<u>Posit Team 2022</u>). Figures were made using the packages *ggplot2* (<u>Wickham 2016</u>) and *patchwork* (<u>Pedersen 2020</u>). In addition, packages *sf* (<u>Pebesma 2018</u>), *spdep* (<u>Bivand 2022</u>), *raster* (<u>Hijmans 2022a</u>), *terra* (<u>Hijmans 2022b</u>) and *geodata* (<u>Hijmans et al. 2022</u>) for spatial analyses, as well as *lubridate* (<u>Grolemund and Wickham 2011</u>), *glue* (<u>Hester and Bryan 2022</u>) and *tictoc* (<u>Izrailev 2021</u>) were used. The source code for the entire analysis, from data preparation to statistical validation of metrics and to producing the final manuscript, is available on the <u>GitHub repository</u> for the study.

## 3 Results

### 3.1 Data quantity

Although there has been a general and consistent increase in the number of participating birders and the number of checklists uploaded over the last few years (due to a general trend of eBird growth in India), and these numbers varied considerably between months of a year, the numbers for each individual month also consistently increased year on year across the four-year period (Fig. 2) with very few exceptions. One such exception is a slightly lower participation (number of birders) in the first wave of COVID-19 in April of MY 2019–20 compared with April of MY 2018–19, but this was still associated with more checklists uploaded. Nevertheless, it is clear that eBird data quantity during the pandemic years was not notably anomalous in any month. The decrease in data generation and participation following February is a normal pattern and is not limited to the pandemic years, given the extremely high participation in the global event, Great Backyard Bird Count, in February. In fact, there was even an uncharacteristic increase in data generation in April 2020 (Fig. 2A). Therefore, there is no clear evidence for data quantity having been negatively impacted by the pandemic.

**Figure 2:**
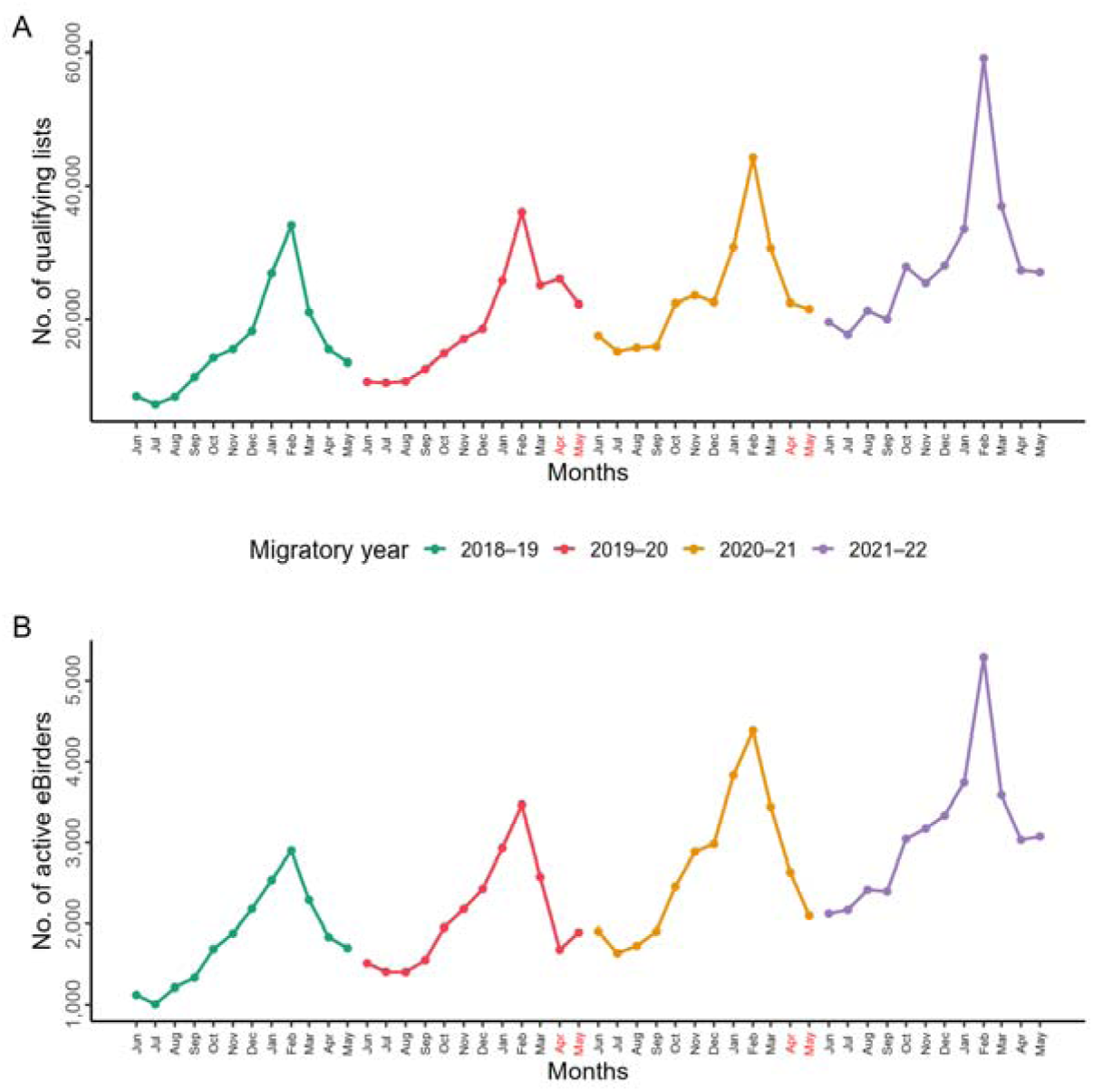
Graphs of change in data quantity over time, in terms of data generation (number of qualifying lists; A) and participation (number of active eBirders; B). Values for the four migratory years are shown in different colours, and the peak pandemic months are shown in red text. Note the peak in data generation and participation in February of each year, which coincides with the global event, Great Backyard Bird Count.

### 3.2 Birder behaviour, urban bias, and spatial coverage

Overall, the characteristics of nationwide eBird data changed predictably during the first wave of the pandemic (April–May, MY 2019–20; red ribbon in Fig. 3). Interestingly though, these changes were short-lived and the metrics returned to their original levels in the period of June 2020–March 2021 (blue ribbon). In fact, changes were minimal even during the second peak pandemic period (April–May, MY 2020–21). Some individual states also showed an indication of a similar pattern, though uncertainty in estimates was high (Fig. 4).

**Figure 3:**
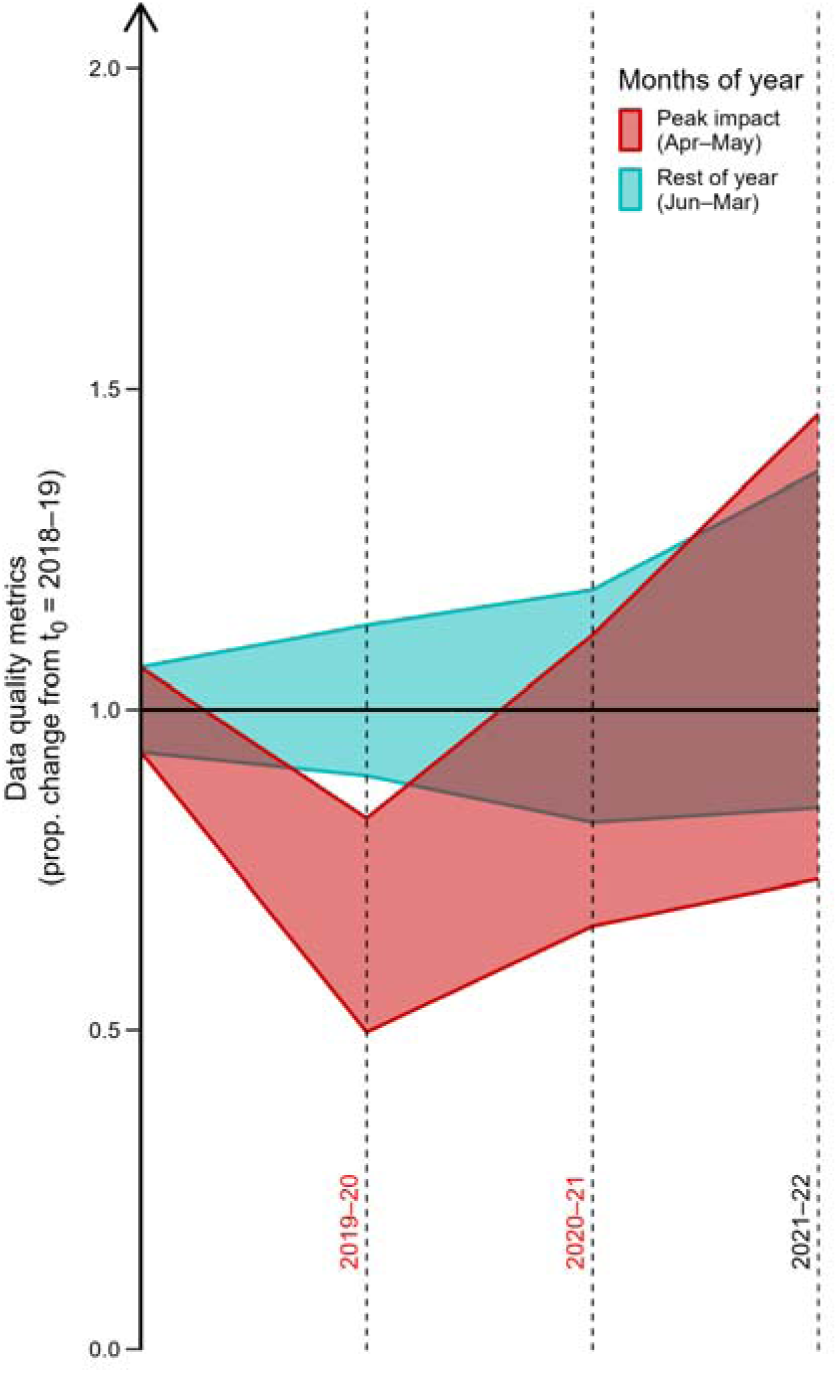
Proportional change in data quality metrics per year and month category, relative to values in MY 2018–19 (represented by the y = 1 line). The four years of interest are on the X-axis, with the pandemic years labelled in red text. Coloured ribbons represent 95% confidence intervals, obtained from propagated error, for the two sets of months. Note that means have not been depicted.

**Figure 4:**
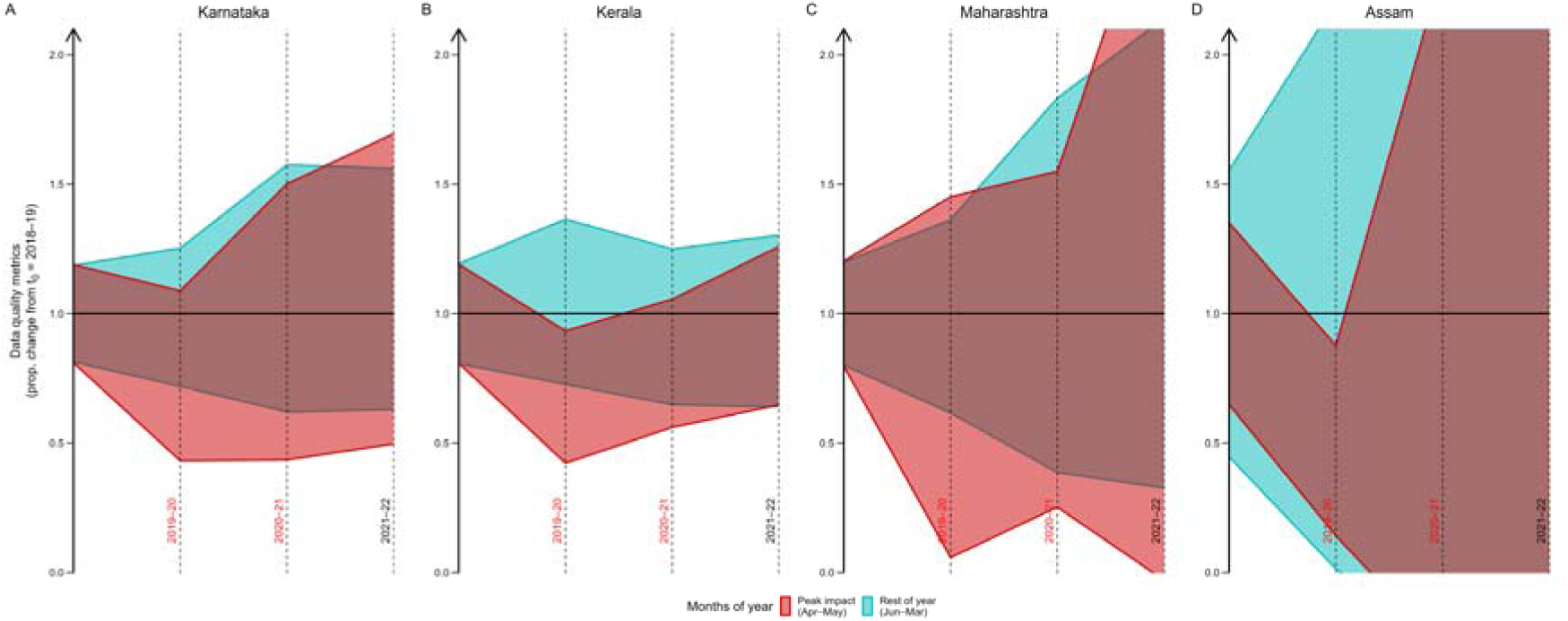
Proportional change in data quality metrics per year and month category, relative to values in MY 2018–19 (represented by the y = 1 line). This is visualised for the four states of Karnataka (A), Kerala (B), Maharashtra (C), and Assam (D). The four years of interest are on the X-axis, with the pandemic years labelled in red text. Coloured ribbons represent 95% confidence intervals, obtained from propagated error, for the two sets of months. Note that means have not been depicted.

Across both spatial scales, although data characteristics in MY 2021–22 are indistinguishable from those in MY 2018–19, they have much higher variability, but note the magnitude of growth in eBird data generation and participation over this period (Fig. 2A, B).

Birders behaved as might be expected during the peak pandemic months, especially during April–May of MY 2019–20. There were prominent declines in birding instances (as defined by individual checklists) with groups of birders (Supplementary Material Fig. S1A, Table S1) and in eBird hotspots (Supplementary Material Fig. S1C, Table S3). Birders seemed to switch to a more sedentary birding style in the peak period, preferring to make stationary checklists over travelling ones (Supplementary Material Fig. S1D, Table S4), showing higher site fidelity, i.e., travelling to fewer spatial units than usual (Supplementary Material Fig. S1B, Table S2), and also being more confined to urban areas than usual (Supplementary Material Fig. S3A, Table S8). Although birders collectively covered slightly fewer grid cells during April of MY 2019–20 (Supplementary Material Fig. S3B), this did not translate to the larger temporal scale of years (Supplementary Material Table S9). In fact, coverage only increased in every successive month since and over the years (Supplementary Material Fig. S3B).

Aside from these suggestions of direct impacts of the pandemic on birder behaviour and data characteristics—those metrics that changed during the peak months and reverted to “normal” soon after—there were also other changes that seem to reflect broader and more general changes over time. For instance, although the pandemic-induced shift to a more sedentary birding style did result in shorter distances covered by the minority birders that made travelling checklists (Supplementary Material Fig. S4A, Table S5), there is also a general trend towards shorter travelling lists that started before the pandemic and has persisted even after. Similarly, birders started making shorter duration checklists (Supplementary Material Fig. S4B, Table S6) that reported fewer species (Supplementary Material Fig. S4C, Table S7), both during the peak pandemic months as well as more generally. Temporally, the impact of the pandemic on these aspects seems minimal, as these changes were not restricted to the peak months or even the pandemic years; instead, these broader trends could be related the general growth of birding and eBirding in India and a potential accompanying change in birding attitudes (see Discussion), but note that decline in checklist-level effort does not imply decline in overall effort.

### 3.3 Spatial spread

Both transition periods (BEF–DUR and DUR–AFT) witnessed an overall positive raw change in birding effort, as defined by number of checklists, across districts (Fig. 5A, Fig. 6A). This was true also for grid coverage, where although the net change in DUR–AFT was lower than in BEF–DUR (due to more districts showing lower coverage in DUR–AFT; Fig. 5B), it was still not negative across districts (Fig. 6C). There were very few major raw changes, and these were not clustered in space. Proportional changes (Fig. 7) included a few major declines (bright yellow) in both birding effort and grid coverage in BEF–DUR and AFT–DUR, but these were restricted to a few districts in central and northern India and there was little overlap between the two transitions. Moreover, there were also similar numbers of districts showing prominent declines in the opposite direction, i.e., in DUR–BEF and DUR– AFT. More districts showed major declines when going backwards in time (DUR–BEF and AFT–DUR) than forwards. Several districts were covered during the pandemic that were not covered before or after.

**Figure 5:**
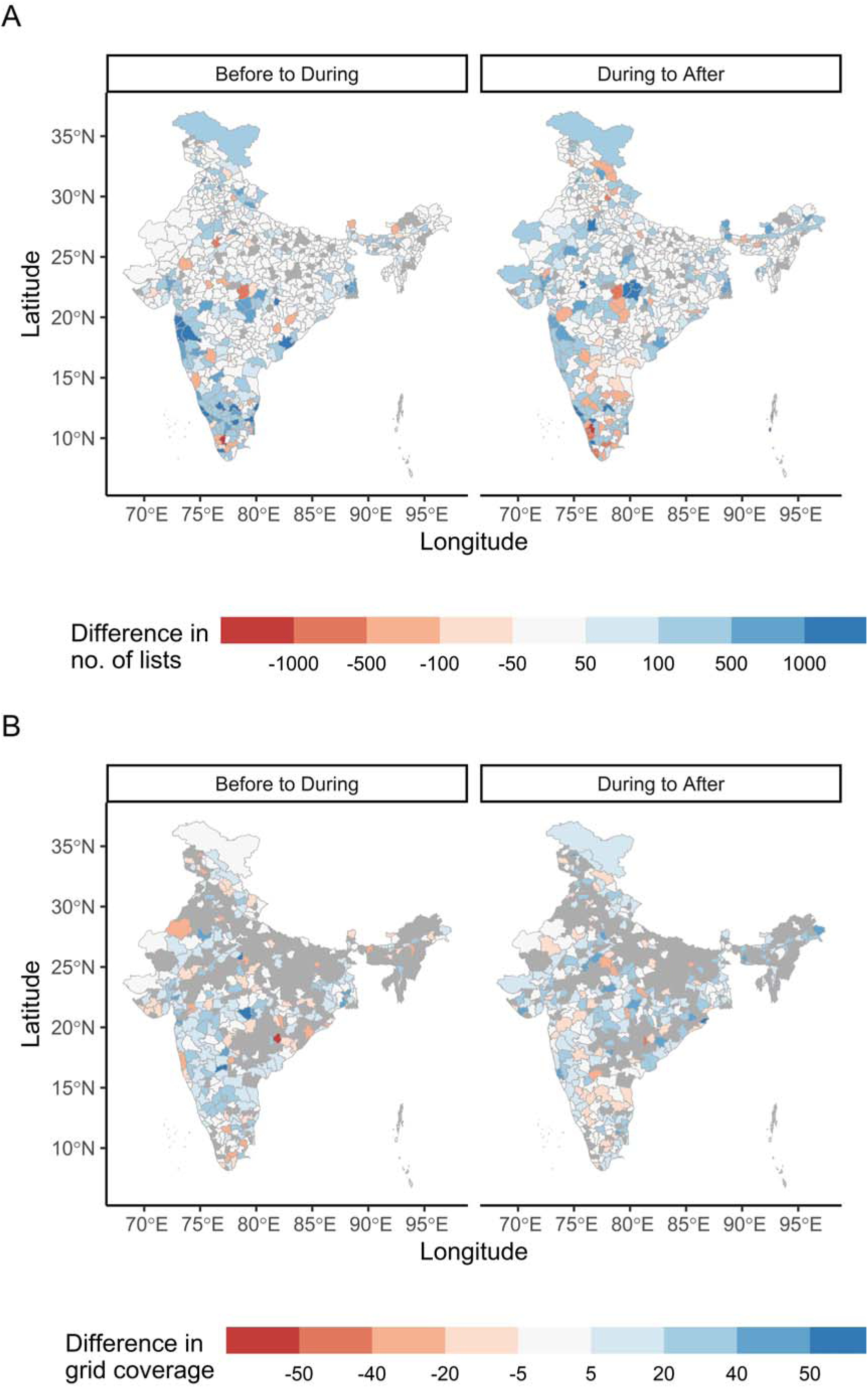
Raw change in number of lists (A) and thresholded percentage grid coverage (B) per district from before to during and from during to after the pandemic. Districts are shaded based on the simple difference in values between the two periods from highly negative (red) to highly positive (blue). Note the absence of spatial clustering among districts with extreme changes, especially declines. Grey districts are those that did not meet the criteria for inclusion in the analysis.

**Figure 6:**
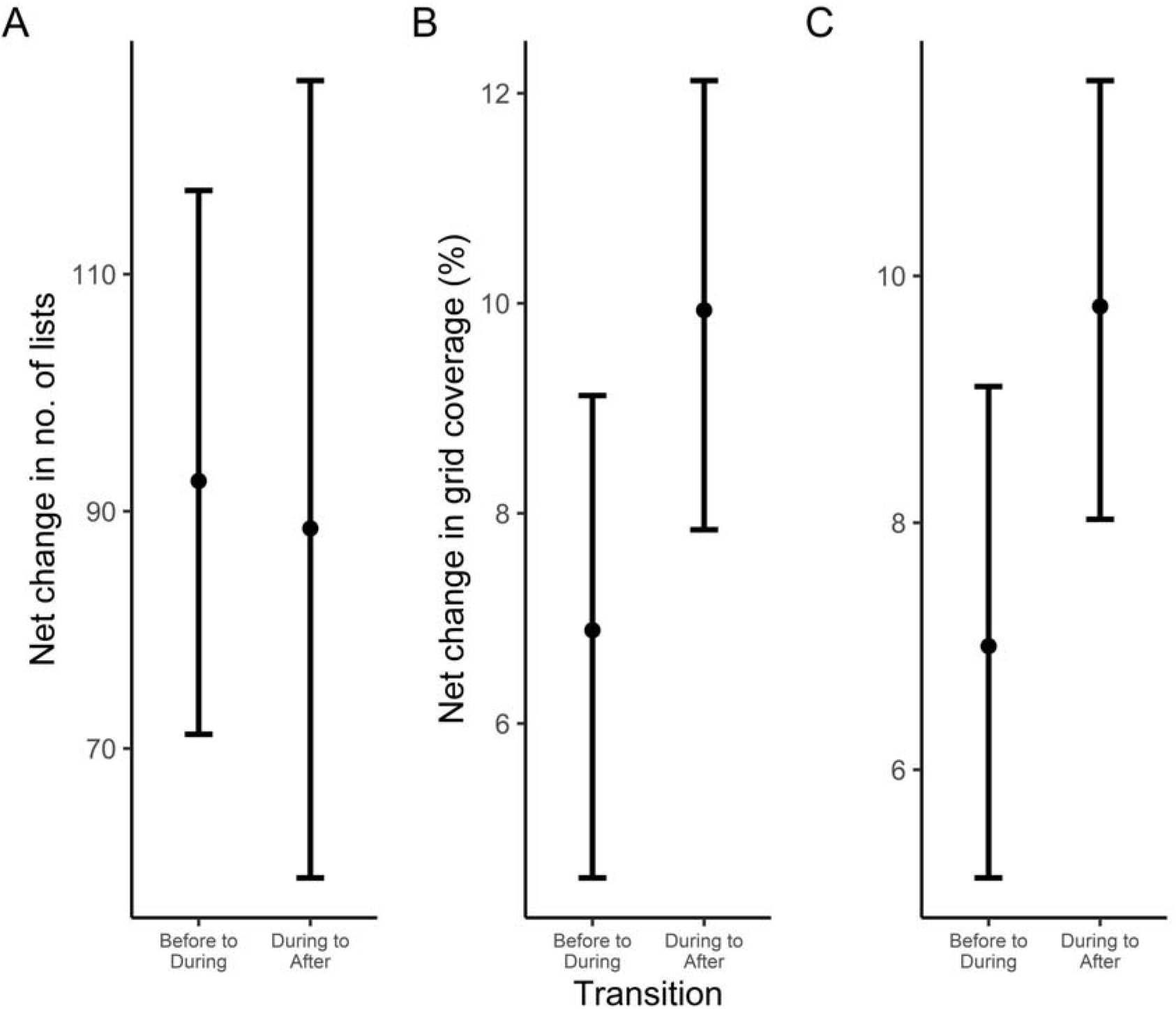
Net change across districts in number of lists (A), percentage grid coverage with no threshold (B) and thresholded percentage grid coverage (C) from before to during (BEF– DUR) and from during to after (DUR–AFT) the pandemic.

**Figure 7:**
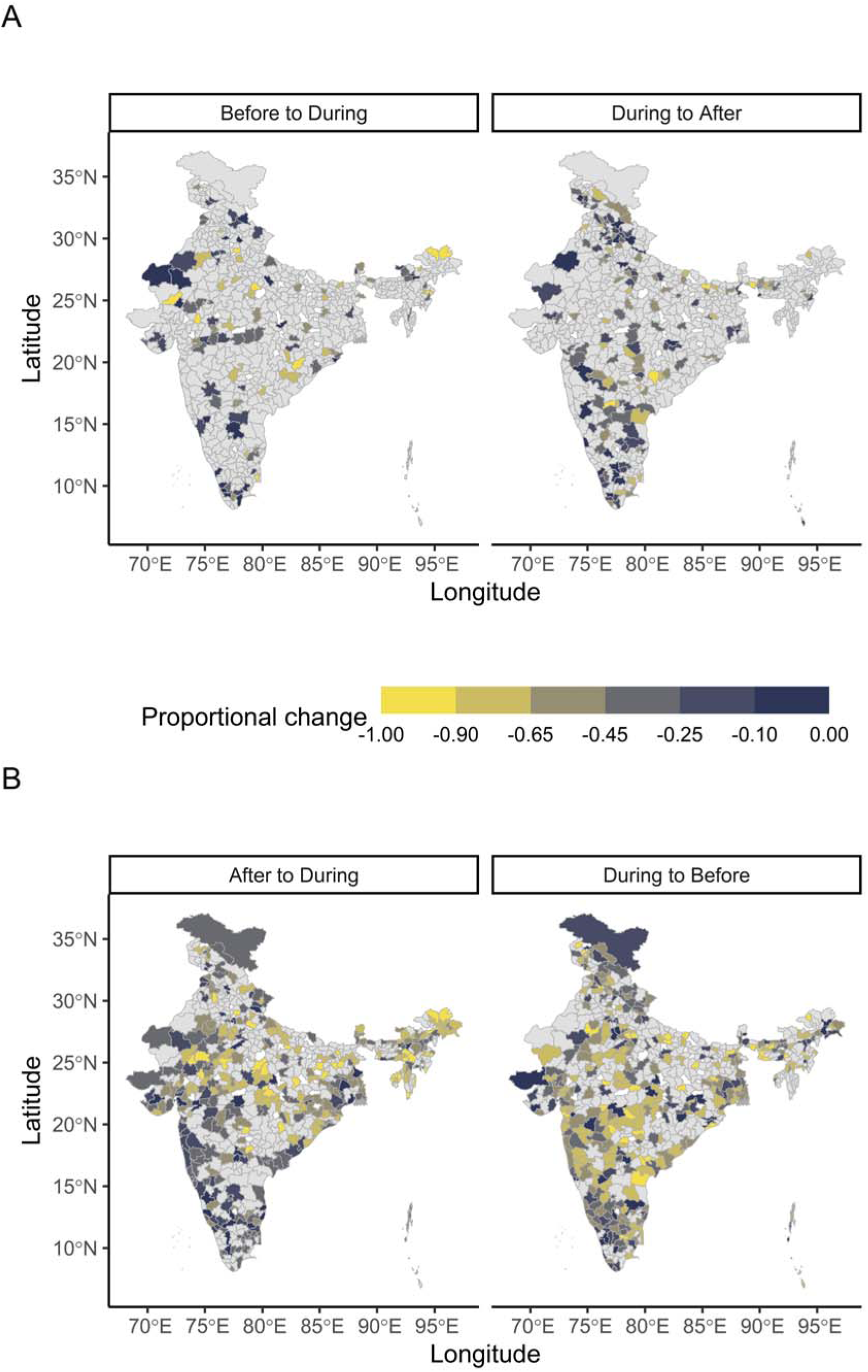
Proportional changes in number of lists per district in forward (top row) and backward (bottom row) transitions. Districts with positive values are shown in grey; declines are shaded based on the magnitude of proportional change from low (dark blue) to high (yellow). Note the absence of spatial clustering among districts with major declines over time (top row). White districts are those that did not meet the criteria for inclusion in the analysis.

### 3.4 Scale of patterns

Pandemic-induced changes were largely transient: they were limited to the peak months of the pandemic, and these unsurprising fine-scale changes did not translate over to the larger scale of years. Among these short-term changes to birder behaviour, the decline in group birding lasted the longest, extending even to a few months after the first peak pandemic period (Supplementary Material Fig. S1A). Notably, birders recovered much quicker in the second year than in the first (Fig. 3). These patterns of birder behaviour—particularly the transience of changes—were largely consistent across different states (Fig. 4) with mostly only minor variations (e.g., Supplementary Material Fig. S2). Major differences were rare, but one example is the complete absence of urban bias in Kerala. Interestingly, Kerala was also the only one of the four states to perfectly mirror the national-level pattern of consistent and unambiguous decline in data quality metrics during the peak months of MY 2019–20 (Fig. 4B); others showed high variability within the state, with some instances even of increases (e.g., Maharashtra). Overall, across these spatial units, data characteristics therefore remained similar in the years before, during and after the pandemic.

## 4 Discussion

### 4.1 Birders and the pandemic: early predictability but innate resilience

The months of April and May in MY 2019–20 and MY 2020–21 were the peak periods of the pandemic in India and involved some of the most stringent restrictions and government regulations (e.g., nationwide lockdown) in order to curb the spread of the disease (<u>Gettleman and Schultz 2020</u>, <u>BBC News 2020</u>, <u>The Economic Times 2020</u>). Accordingly, most changes to eBird data were also concentrated in these peak pandemic months. Unsurprisingly, birders became more isolated during these periods (especially in the first lockdown in April–May 2020), avoiding travel, groups, and public spaces. They switched to a more sedentary birding style, minimising movement and travel, showing higher site fidelity and being more confined to urban areas than usual. Naturally, this also resulted in a slight decline in spatial coverage across the country. Birders also made shorter checklists that reported fewer species. Thus, birders adapted their behaviour during the first peak pandemic period in accordance with the various official guidelines to prevent and minimise the spread of the disease.

As restrictions were gradually lifted in the following months, birder behaviour also changed back to normal, until the peak months of the second pandemic year. Despite this second wave of the pandemic and lockdowns being more devastating than the first in terms of COVID-19 cases and fatalities (<u>Worldometer 2023</u>), changes in birder behaviour (such as isolation) were relatively muted and also recovered more quickly this time. The staggered recovery in the first year and the almost immediate recovery in the second suggest that the novelty of the disease played an important role in changing birder behaviour, and also point to the innate resilience of birders to perturbation. As a result of this, all metrics of eBird data characteristics (i.e., birder behaviour, urban bias, spatial coverage, spatial spread) remained unaffected at the yearly level (see coefficients for years in Tables 1–9).

### 4.2 Evolution of eBirding in India

We found that eBird data quantity has been consistently increasing over the years, in terms of both participation and data generation. This was true even during the peak pandemic periods, despite the various restrictions on movement and social interaction. Such increases in data quantity have been observed elsewhere, both in eBird (<u>Hochachka et al. 2021</u>) and in other CS programmes like iNaturalist (<u>Crimmins et al. 2021</u>), but in our case we found little evidence for this increase being directly linked to the pandemic. Instead, our results suggest a consistent and long-term growth of eBirding in India over the years, which has also resulted in an increase in overall spatial coverage of eBird data.

Furthermore, this steady growth seems to have been accompanied by shifts in India’s birding and eBirding culture, but note that specifics vary considerably depending on the region and scale under consideration. Some of the observed trends in birder behaviour, although exaggerated during the peak pandemic months, have existed since MY 2018–19 (i.e., started even before the pandemic) and have persisted much later after the pandemic has passed. In particular, checklist-level effort has been consistently lower in terms of duration, distance and length (number of species reported). This suggests a gradual evolution towards shorter birding sessions (checklists) and finer scale data generation, even if overall birding effort is not necessarily lower. Of these three metrics, it is worth noting that list length appears to be more resilient than distance and duration to perturbations like the pandemic and to gradual changes in birder behaviour, adding to the growing evidence that list length is the more robust metric of effort in analyses of eBird data (<u>Viswanathan n.d.</u>).

### 4.3 Utility of eBird data from the pandemic

The appreciation and use of large-scale, crowdsourced, open access data such as from CS programmes have been growing in recent years, and came into the spotlight during the unprecedented and disruptive times of the COVID-19 pandemic which drastically altered human circumstances worldwide. Most studies linking the pandemic and CS data have focused on changes at the organismal level (e.g., <u>Warrington et al. 2022</u>), exploring the phenomenon of anthropause (<u>Rutz et al. 2020</u>) to better understand the influence of anthropogenic factors on organisms (<u>Bates et al. 2020</u>).

In contrast, there is a paucity of studies looking at the effects of the pandemic on the very characteristics of CS data (i.e., observer/human level changes). We expected changes to the nature of eBird data during the pandemic due to behavioural changes in the birding community, caused directly by the various restrictions and indirectly by lifestyle changes. What we found is a steady increase in data quantity, and certain expected but short-lived changes to birder behaviour. Our findings echo the interpretation of Hochachka et al. (<u>2021</u>) of their own results: compared with the magnitude of change brought by the pandemic, its impacts on birder behaviour and thereby eBird data are not substantial. While our results pertain to birders and eBird, there is precedent to find similar trends in other CS programmes as well. We posit that this is due to a stronger urge during the pandemic (and perhaps other challenging circumstances) for people to connect with nature (<u>Marsh et al. 2021</u>), as well as an inherent resilience in birders that buffers them from radical changes to their birding lifestyle. In other, more aphoristic words, birders will be birders.

Thus, we find that the pandemic has not caused major changes in the utility of eBird data, particularly for applications such as abundance trend estimation which are relatively more vulnerable to changes in data quality (than, say, range estimation). Our results suggest that eBird data from the pandemic remain immensely valuable, as well as highlight certain key points regarding using this data and making inferences from it.

#### 4.3.1 Importance of scale

Hochachka et al. (<u>2021</u>) emphasised the importance of considering spatial scale, by showing the presence of high degrees of regional variability in the effects of the pandemic. However, they only considered a single month across years, and so failed to capture variability across the temporal scale. Our findings highlight that considerations of both spatial and temporal scales are crucial in assessing the impacts of a major event like the COVID-19 pandemic.

At the fine scale of months and states (or even smaller like districts), there were indeed clear and expected changes in birder behaviour and, consequently, to the nature of the data. These fine-scale changes were scattered throughout the country and not concentrated in one or a few specific regions. Notably, most changes showed high variability (Fig. 4) and thus averaged out across different states (spatial nuance, due to differences in myriad factors that are difficult to predict, such as social and political scenarios), and reverted to regular levels after the lockdown months (temporal transience).

As a result, there were very few notable nationwide changes across the four years (i.e., large spatial and temporal scale). Most analyses of long-term abundance trends, distributions or migratory patterns using eBird data are carried out at such large scales, and hence can continue to be performed.

#### 4.3.2 Importance of effort

A detailed understanding of how birding effort in India has changed over the years requires a longer-term analysis of metrics of effort, starting from the inception of eBird in India. While this is beyond the scope of our study, we did find suggestions of such a change, neither caused by nor restricted to the COVID-19 pandemic but instead representing a consistent directional change as Indian birding evolved and is likely to stay.

Therefore, it becomes important to appropriately account for this variability in effort in any analysis spanning multiple years. Viswanathan (<u>n.d.</u>) showed that within individual geographies where species detections and accumulation can be assumed to be relatively constant, list length has a tighter relationship with the data generation process than distance and duration. Our results suggest that list length is also more resilient to sudden perturbations as well as to year-to-year variability.

#### 4.3.3 Guidelines for using pandemic eBird data from India

We outline some guidelines for making use of the tremendous potential in CS programmes like eBird even during the pandemic. When formulating a question to be answered using this data, the researcher must first consider what scale (both spatial and temporal) is appropriate for the question. Questions of larger scale can be tackled directly, whereas fine-scale questions will benefit from a deep-dive into artefacts in the raw data that can produce misleading patterns. In the latter cases, the researcher is recommended to conduct analyses of data characteristics (similar to the ones in this study) in order to first assess the suitability of the data before attempting to answer the question. Variability in sampling effort is likely to be an issue at smaller scales, but its effect can be minimised by including effort variables such as list length as well as by using methods of spatial subsampling in the analyses. In both cases (large and small scales), it is important to ensure that analyses being conducted are locally appropriate: after identifying the relevant scale and effort metrics, these should be accounted for appropriately, considering the various idiosyncrasies and nuances that may manifest across space and time.

## 5 Conclusion

We found that birders did indeed behave differently during the pandemic, but its consequences for eBird data characteristics were lower than expected from the impact the pandemic has had on human life. Interestingly, our results suggest that official restrictions introduced during the peak periods of the pandemic, together with its novelty, were most instrumental in changing birder behaviour, while changes in lifestyle were not drastic or long-lasting enough to alter behaviour in the long run. This resilience of birders helped maintain similar data quality levels at large scales, and also led to a steady increase in data quantity. Therefore, the eBird dataset remains valuable, with only certain statistical and scale considerations to be kept in mind.

