## Supplementary Material for "Birdwatchers’ resilience to perturbation in India buffers citizen science from pandemic-induced biases"

### 1 Methods

#### 1.1 Data preparation

##### 1.1.1 Data filters

We standardised the dataset before analysis with the following set of filters. First, we filtered the dataset for “complete checklists”, in which observers report all the birds they see or hear. Next, we filtered the data for our timeline of interest. The migratory cycle in India starts in June of one calendar year and ends in May of the next, and this time frame is thus deemed more appropriate than a calendar year. Therefore, all analyses presented in this study use “migratory year” (MY) as a unit time period.

We then removed data from group accounts, which are eBird accounts with which multiple existing eBirders share their lists to track aggregate summaries. Such accounts are commonly used during specific events or surveys, and need to be removed from analyses of observer-related metrics. Next, we applied a nocturnal filter to remove observations made between 20:00 h and 04:00 h (nocturnal birding is very different from diurnal birding), a pelagic filter to remove observations made from beyond the shoreline of the country, and finally a set of filters to remove observations made with extreme values of birding “effort”: checklists that are too short (less than three minutes long, or reporting three or fewer species without information on duration, or reporting three or fewer species at a rate lower than two species per hour), and travelling checklists that are too fast (following “Traveling” protocol and travelling faster than 20 kmph or covering a distance greater than 50 km). These lists with extreme effort are not considered to be “truly complete” checklists. Finally, in instances of group birding (column “GROUP.IDENTIFIER”) where each participant eBirder owns a copy of the checklist, duplicate checklists were removed by randomly sampling one out of the multiple checklists (column “SAMPLING.EVENT.IDENTIFIER”).

The checklists that passed these criteria are referred to as “qualifying checklists”.

##### 1.1.2 Urban or rural

We used remote sensing data to divide the country into urban and non-urban (rural) areas. For this, we downloaded the MODIS Land Cover Type Product (MCD12Q1) that includes several science datasets (SDSs) mapping global land cover at a 500 m resolution. Although this comes at an annual time step, we used data from the single time point, 31 December 2021. The tiled MODIS data was downloaded using the “getModis()” function in the *luna* package and saved as individual .tif files, which were then merged into a single raster. After cropping and masking to the country’s administrative boundary shapefile, the MODIS land cover categories in the raster were reclassified: category 13, “urban and built-up lands” was considered “urban”, and all other categories were considered “rural” (technically “non-urban”). We didn’t use the MODIS data at 500 m × 500 m resolution because the location information in eBird data, despite being precise, is not very accurate at small grain sizes (due to the very nature of eBird data where sampling often occurs over a distance). So, finally we aggregated the raster to a resolution of 2 km $\times$ 2 km which was deemed more appropriate, using a 25% threshold for classification as urban, i.e., a 2 km $\times$ 2 km cell was classified as urban if four or more of the 16 500 m $\times$ 500 m cells being aggregated were urban. We then linked each qualifying checklist in the dataset to this modified MODIS raster, using its associated latitude-longitude information, and thus classified each checklist as urban or rural.

#### 1.2 Spatial spread

For analysing spatial spread of the data, we decided to focus on districts as the spatial unit instead of 0.225° $\times$ 0.225° grid cells as in other metrics. This is because birding patterns in general are not uniformly and consistently spread across the country, so the fine scale of grid cells would produce too much noise. Here, districts provide a nice compromise: they are large enough to track change with less noise, while still being small enough to inform us about local patterns. We ensured that a single grid cell was linked with only one district, using random sampling from the list of districts it covered. The temporal unit of focus was the categorical COVID time period (“BEF”, “DUR” or “AFT”) and not individual months. Only those districts that were “covered” at least once in the study period, i.e., had at least one qualifying checklist in migratory years 2018–2021, were considered in the analyses.

Our aim was to understand whether (and if so, how) the way birding patterns were spread across the country changed during the pandemic years. The two patterns of interest were change in birding effort (number of lists) and change in grid coverage (proportion of total grid cells having birding activity) from one year to the next. The former tells us about the intensity of birding while the latter shows us its breadth or completeness. Here, the changes in metrics rather than the metrics themselves inform us about consistency/variability across years, and would better highlight any anomalies brought by the pandemic. In the case of the period “DUR”, both metrics were averaged across the two migratory years it comprised, 2019–20 and 2020–21.

The grid coverage metric considered only those districts comprising at least five grid cells after the randomised assignment of grid cells to districts. Although this meant that certain districts which actually cover five or more grid cells might have been disqualified due to the random assignment, the converse is also true so some districts which in reality do not cover five or more grid cells might have qualified. We assume the two cases average out over the entire country, so it is not a statistical concern. However, grid coverage was further analysed in two conditions: one with no threshold for grid cell coverage, and one where a cell was considered “covered” only if it had five or more qualifying checklists per migratory year. The latter condition, by removing outlier cases, would produce clearer patterns with less noise.

Due to the difficulty in quantitatively ascertaining change in spatial spread across such a large area with huge variability, we devised a suite of three analyses which, put together, would provide a clear qualitative picture of change in spatial spread/clustering of the metric.

##### 1.2.1 Spatial visualisation of raw change

Raw change in a metric $a$ from period $x$ to period $y$ was calculated as the simple difference of their values $\left( a_{y}-a_{x} \right)$. It was not calculated for cases where both $a_{x}$ and $a_{y}$ were zero. We plotted the raw change per district from one COVID period to another on a map of India, colouring the districts based on magnitude of change. Here we were concerned with two things: firstly, whether the majority of districts showed a positive or a negative change; second, how many districts showed high magnitude of change, and whether or not these were clustered in space. If most districts showed a negative change in birding effort or grid coverage from BEF to DUR, this would translate to an effect of the pandemic on the quantity and quality of data collected during the pandemic. The higher the magnitude of change, the more worrisome the change would be, and if these high magnitude changes were spatially clustered, this would result in a high spatial bias during the pandemic years, as opposed to if they were spread out across different regions of the country.

##### 1.2.2 Net raw change across the country

This is a more quantitative method of analysing whether the majority of districts showed a positive or a negative change. We estimated mean raw change and its 95% CI, by bootstrapping across all the districts. If the net change from DUR to AFT was significantly (no overlap of CIs) higher than from BEF to DUR, this would suggest that birding effort or grid coverage was subdued during the pandemic.

##### 1.2.3 Spatial visualisation of proportional change in two temporal directions

Proportional change in a metric $a$ from period $x$ to period $y$ was calculated as $\left( a_{y}-a_{x} \right)/a_{x}$. It was not calculated for cases where both $a_{x}$ and $a_{y}$ were zero. If only $a_{x}$ was zero, a constant $k=1$ was added to both $a_{x}$ and $a_{y}$ in order to avoid zero in the denominator. Moreover, for the birding effort metric, proportional change was not calculated if the first period in the transition had less than 10 qualifying checklists ($a_{x}<10$). Otherwise, since the metric has a fixed lower limit but not an upper limit, this would result in very high values of proportional change simply due to the low overall birding.

We visualised only the negative values of proportional change, since a general increase in birding effort or grid coverage is not worrisome and is in fact even expected. Here, the values were coloured based on magnitude and once again the high magnitude changes are of interest, but we are also interested in transitions in both directions. For example, if the same districts with high magnitudes of negative proportional change from BEF to DUR have similar values also from AFT to DUR, this would suggest a clear and direct impact of the pandemic in lower birding effort or grid coverage.

#### 1.3 Model specifications

GLMMs were fitted with the following specification, with the random effect changing as appropriate (see Section 2.2.6):

$$metric\sim year+year:month+\left( 1/observer \right)$$

$$metric\sim year+year:month+\left( 1/cell \right)$$

where $year$ represents the migratory years. Where GLMMs were not appropriate, LMMs were fitted with the following specification:

$$log\left( metric \right)\sim year+year:month+\left( 1/observer \right)$$

### 2 Figures

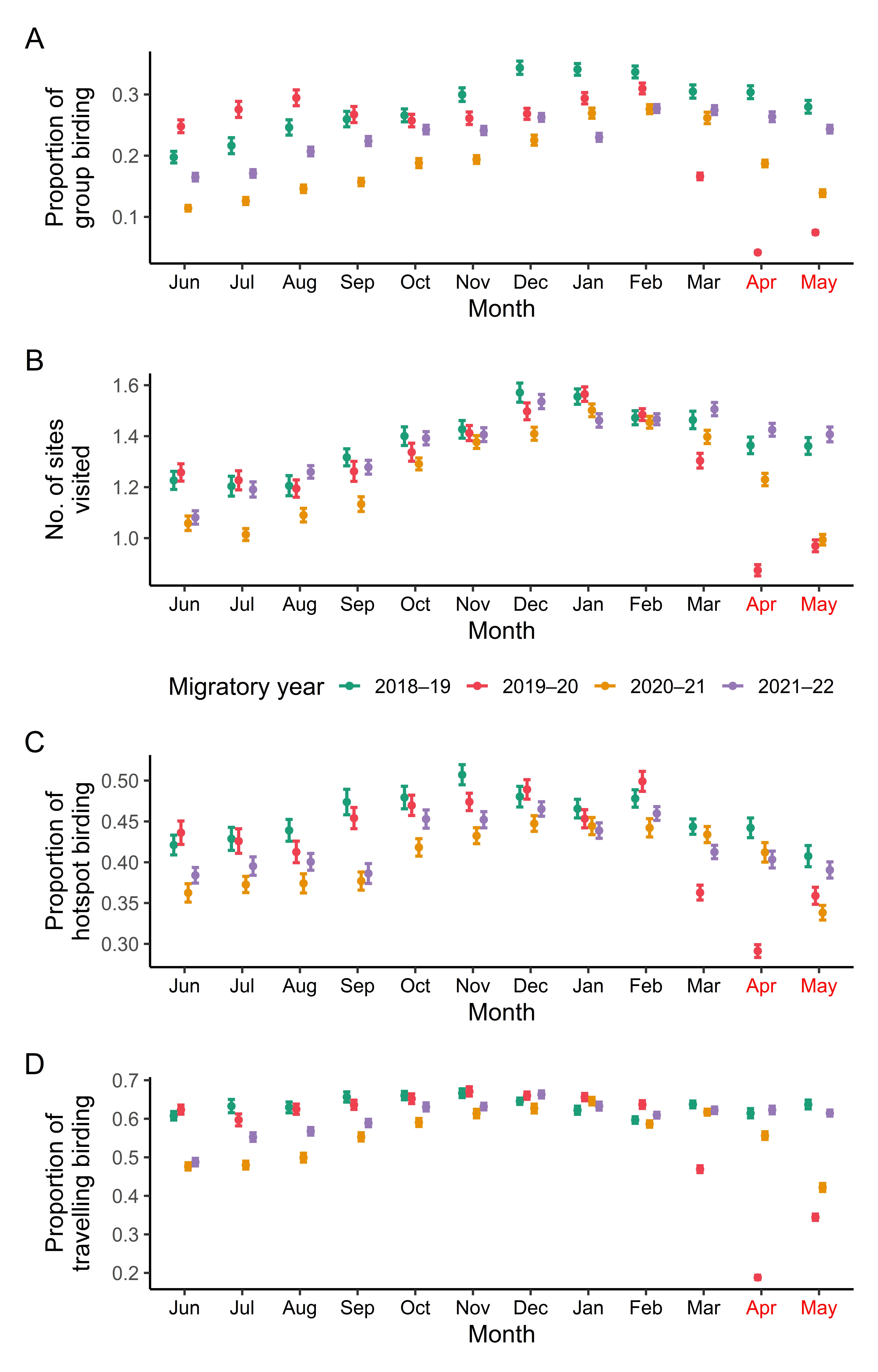

Figure S1: Graphs of observer-level group birding (A) and inverse site fidelity (B), hotspot birding (C) and travelling protocol birding (D) at the national level. Points represent means while error bars show 95% confidence intervals. Values for the four migratory years are shown in different colours, and the peak pandemic months are shown in red text.

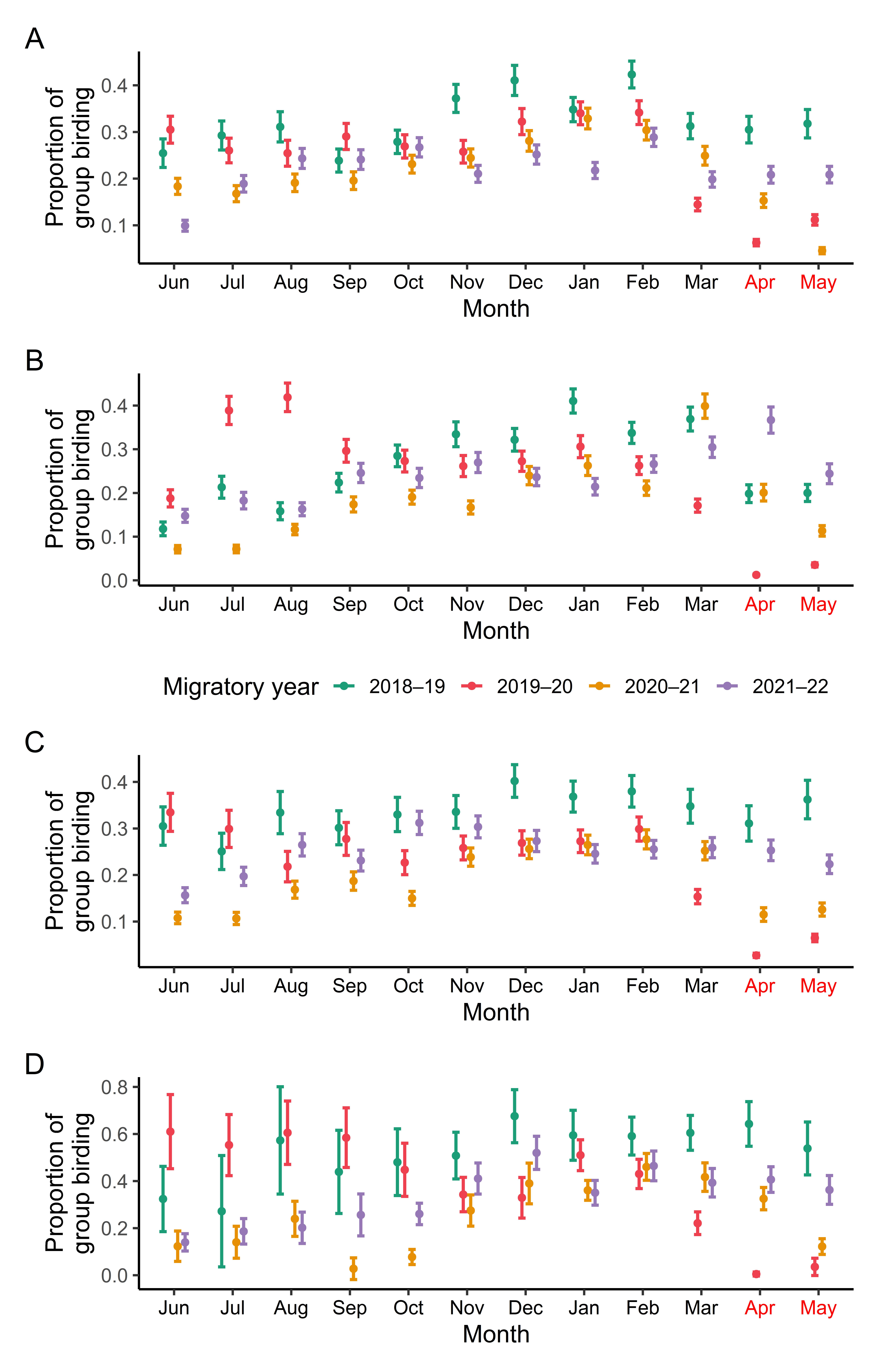

Figure S2: Graphs of observer-level group birding for the four states of Karnataka (A), Kerala (B), Maharashtra (C), and Assam (D). Points represent means while error bars show 95% confidence intervals. Values for the four migratory years are shown in different colours, and the peak pandemic months are shown in red text.

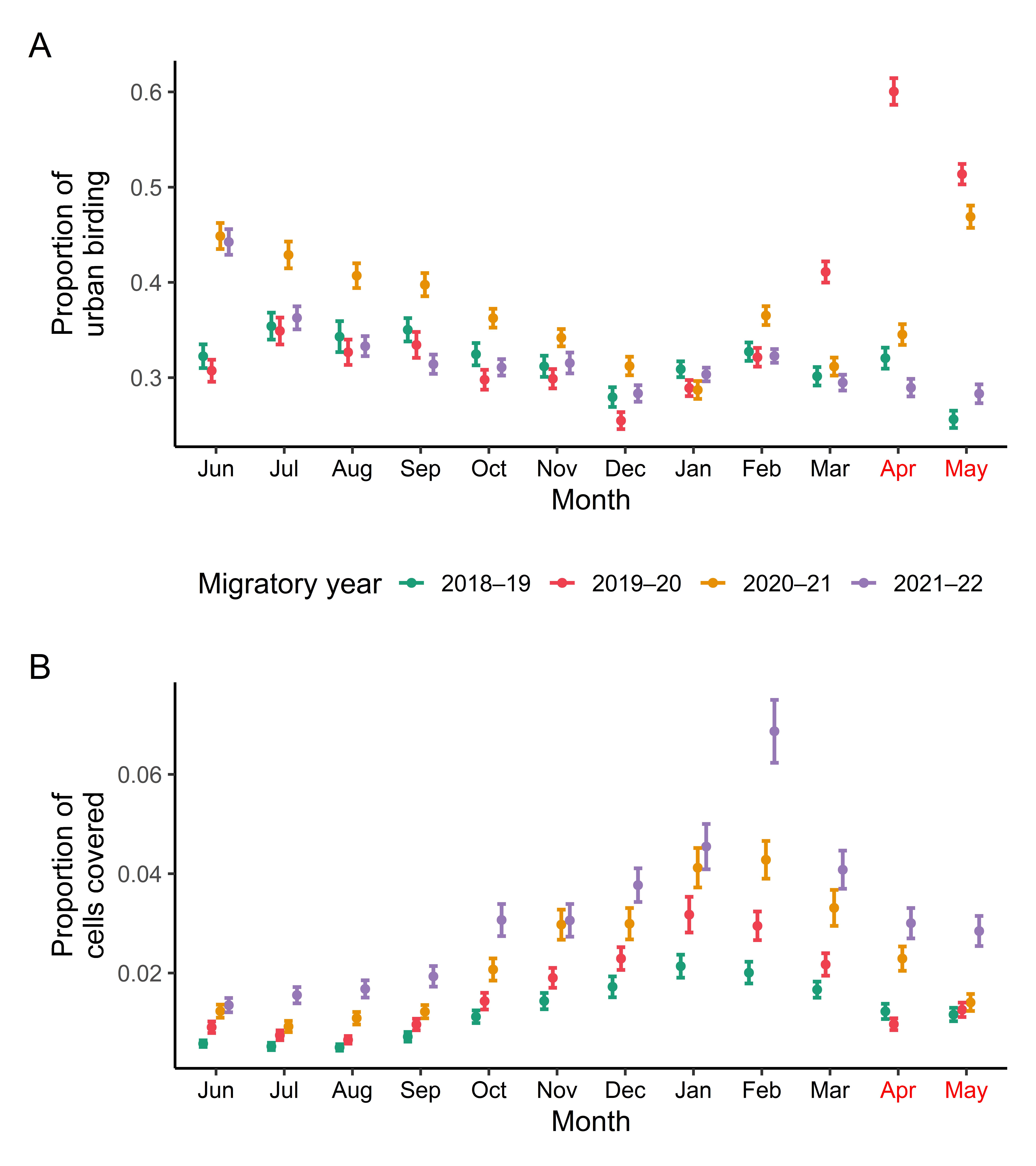

Figure S3: Graphs of urban bias (A) and spatial coverage (B) at the national level. Points represent means while error bars show 95% confidence intervals. Values for the four migratory years are shown in different colours, and the peak pandemic months are shown in red text.

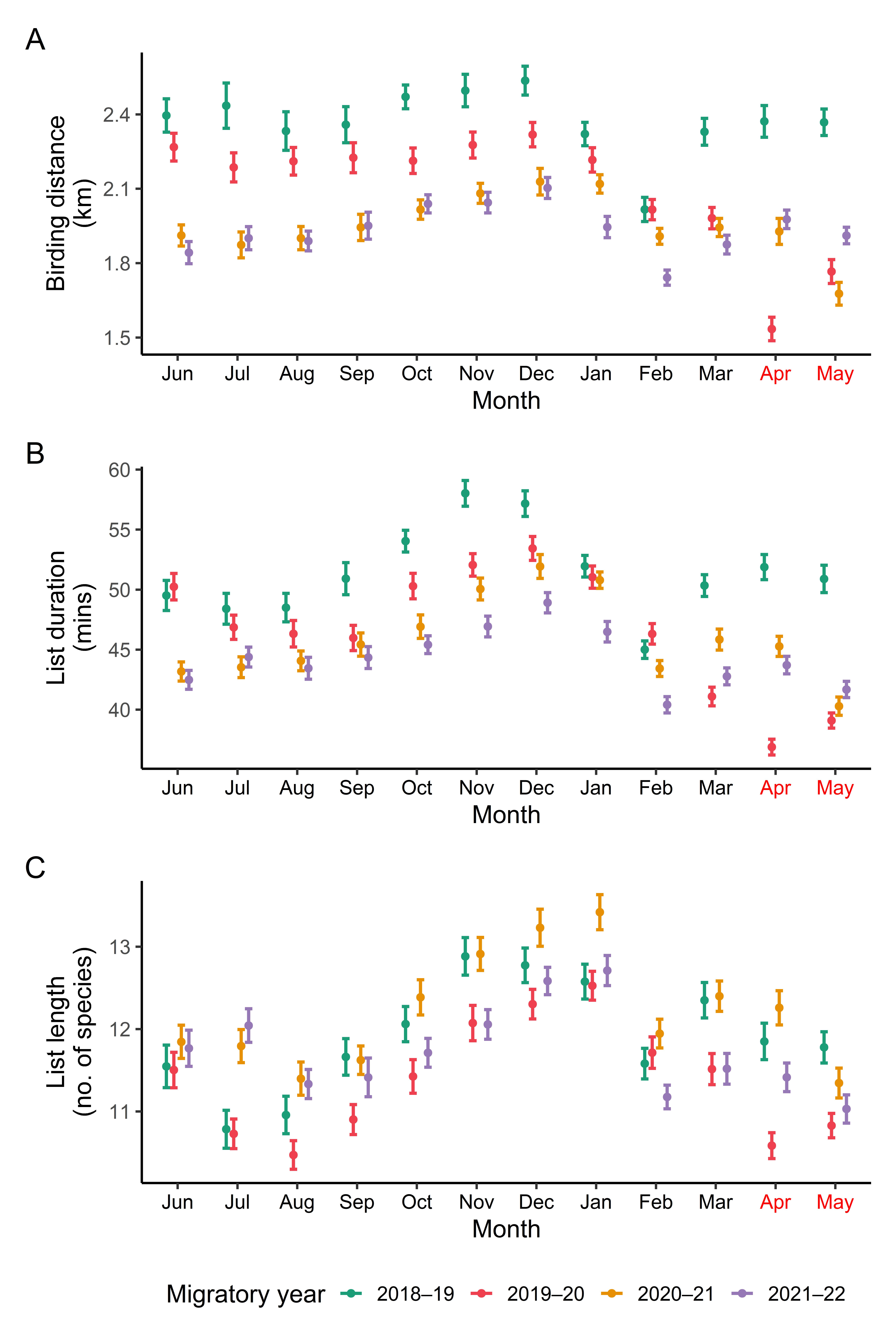

Figure S4: Graphs of birding distance, list duration and list length at the national level. Points represent means while error bars show 95% confidence intervals. Values for the four migratory years are shown in different colours, and the peak pandemic months are shown in red text.

### 3 Tables

Table S1: Model summary for group birding at the national level. Numerical years refer to "migratory years". Random effect variance of Observer ID is 2.515. See model specifications described earlier.

| Fixed effect | Estimate | Standard error |
| --- | --- | --- |
| (Intercept) | -1.119 | 0.022 |
| 2019 | -0.182 | 0.023 |
| 2020 | -0.287 | 0.023 |
| 2021 | -0.473 | 0.023 |
| 2018:February | -0.018 | 0.021 |
| 2019:February | 0.057 | 0.022 |
| 2020:February | 0.025 | 0.021 |
| 2021:February | 0.215 | 0.021 |
| 2018:March | -0.146 | 0.024 |
| 2019:March | -0.649 | 0.027 |
| 2020:March | -0.036 | 0.023 |
| 2021:March | 0.203 | 0.023 |
| 2018:April | -0.144 | 0.027 |
| 2019:April | -2.108 | 0.045 |
| 2020:April | -0.418 | 0.028 |
| 2021:April | 0.162 | 0.024 |
| 2018:May | -0.242 | 0.029 |
| 2019:May | -1.512 | 0.040 |
| 2020:May | -0.750 | 0.032 |
| 2021:May | 0.061 | 0.025 |
| 2018:June | -0.646 | 0.037 |
| 2019:June | -0.199 | 0.033 |
| 2020:June | -0.958 | 0.037 |
| 2021:June | -0.374 | 0.031 |
| 2018:July | -0.542 | 0.038 |
| 2019:July | -0.085 | 0.034 |
| 2020:July | -0.847 | 0.038 |
| 2021:July | -0.331 | 0.032 |
| 2018:August | -0.393 | 0.036 |
| 2019:August | -0.002 | 0.032 |
| 2020:August | -0.688 | 0.034 |
| 2021:August | -0.118 | 0.029 |
| 2018:September | -0.331 | 0.031 |
| 2019:September | -0.113 | 0.031 |
| 2020:September | -0.614 | 0.033 |
| 2021:September | -0.029 | 0.028 |
| 2018:October | -0.302 | 0.028 |
| 2019:October | -0.160 | 0.029 |
| 2020:October | -0.414 | 0.027 |
| 2021:October | 0.061 | 0.025 |
| 2018:November | -0.160 | 0.026 |
| 2019:November | -0.141 | 0.027 |
| 2020:November | -0.380 | 0.026 |
| 2021:November | 0.059 | 0.025 |
| 2018:December | 0.004 | 0.024 |
| 2019:December | -0.114 | 0.026 |
| 2020:December | -0.209 | 0.025 |
| 2021:December | 0.156 | 0.024 |

Table S2: Model summary for site fidelity at the national level. Numerical years refer to "migratory years". Random effect variance of Observer ID is 0.081. See model specifications described earlier.

| Fixed effect | Estimate | Standard error |
| --- | --- | --- |
| (Intercept) | 0.440 | 0.011 |
| 2019 | 0.007 | 0.014 |
| 2020 | -0.031 | 0.014 |
| 2021 | -0.062 | 0.014 |
| 2018:February | -0.055 | 0.015 |
| 2019:February | -0.053 | 0.013 |
| 2020:February | -0.035 | 0.012 |
| 2021:February | 0.005 | 0.011 |
| 2018:March | -0.061 | 0.015 |
| 2019:March | -0.182 | 0.014 |
| 2020:March | -0.071 | 0.012 |
| 2021:March | 0.033 | 0.012 |
| 2018:April | -0.129 | 0.016 |
| 2019:April | -0.579 | 0.016 |
| 2020:April | -0.201 | 0.013 |
| 2021:April | -0.022 | 0.013 |
| 2018:May | -0.132 | 0.017 |
| 2019:May | -0.479 | 0.016 |
| 2020:May | -0.417 | 0.014 |
| 2021:May | -0.037 | 0.013 |
| 2018:June | -0.234 | 0.019 |
| 2019:June | -0.218 | 0.017 |
| 2020:June | -0.351 | 0.015 |
| 2021:June | -0.301 | 0.014 |
| 2018:July | -0.258 | 0.020 |
| 2019:July | -0.243 | 0.017 |
| 2020:July | -0.395 | 0.015 |
| 2021:July | -0.199 | 0.014 |
| 2018:August | -0.253 | 0.019 |
| 2019:August | -0.271 | 0.017 |
| 2020:August | -0.323 | 0.015 |
| 2021:August | -0.148 | 0.014 |
| 2018:September | -0.168 | 0.018 |
| 2019:September | -0.215 | 0.017 |
| 2020:September | -0.282 | 0.015 |
| 2021:September | -0.135 | 0.014 |
| 2018:October | -0.105 | 0.017 |
| 2019:October | -0.154 | 0.015 |
| 2020:October | -0.153 | 0.014 |
| 2021:October | -0.046 | 0.013 |
| 2018:November | -0.085 | 0.016 |
| 2019:November | -0.100 | 0.015 |
| 2020:November | -0.089 | 0.013 |
| 2021:November | -0.037 | 0.013 |
| 2018:December | 0.012 | 0.016 |
| 2019:December | -0.043 | 0.014 |
| 2020:December | -0.064 | 0.013 |
| 2021:December | 0.049 | 0.013 |

Table S3: Model summary for hotspot birding at the national level. Numerical years refer to "migratory years". Random effect variance of Observer ID is 1.858. See model specifications described earlier.

| Fixed effect | Estimate | Standard error |
| --- | --- | --- |
| (Intercept) | -0.599 | 0.018 |
| 2019 | -0.038 | 0.018 |
| 2020 | -0.063 | 0.017 |
| 2021 | -0.082 | 0.017 |
| 2018:February | 0.038 | 0.017 |
| 2019:February | 0.142 | 0.017 |
| 2020:February | -0.006 | 0.015 |
| 2021:February | 0.063 | 0.014 |
| 2018:March | -0.070 | 0.019 |
| 2019:March | -0.288 | 0.018 |
| 2020:March | -0.034 | 0.016 |
| 2021:March | -0.083 | 0.015 |
| 2018:April | -0.069 | 0.021 |
| 2019:April | -0.561 | 0.019 |
| 2020:April | -0.102 | 0.017 |
| 2021:April | -0.113 | 0.016 |
| 2018:May | -0.184 | 0.021 |
| 2019:May | -0.304 | 0.019 |
| 2020:May | -0.353 | 0.018 |
| 2021:May | -0.155 | 0.016 |
| 2018:June | -0.136 | 0.025 |
| 2019:June | -0.046 | 0.022 |
| 2020:June | -0.266 | 0.019 |
| 2021:June | -0.176 | 0.018 |
| 2018:July | -0.121 | 0.025 |
| 2019:July | -0.081 | 0.023 |
| 2020:July | -0.234 | 0.020 |
| 2021:July | -0.133 | 0.018 |
| 2018:August | -0.076 | 0.025 |
| 2019:August | -0.124 | 0.023 |
| 2020:August | -0.231 | 0.020 |
| 2021:August | -0.124 | 0.018 |
| 2018:September | 0.021 | 0.022 |
| 2019:September | 0.005 | 0.021 |
| 2020:September | -0.216 | 0.019 |
| 2021:September | -0.171 | 0.018 |
| 2018:October | 0.037 | 0.021 |
| 2019:October | 0.049 | 0.020 |
| 2020:October | -0.085 | 0.017 |
| 2021:October | 0.046 | 0.016 |
| 2018:November | 0.122 | 0.020 |
| 2019:November | 0.064 | 0.019 |
| 2020:November | -0.038 | 0.017 |
| 2021:November | 0.043 | 0.016 |
| 2018:December | 0.043 | 0.019 |
| 2019:December | 0.113 | 0.019 |
| 2020:December | 0.013 | 0.017 |
| 2021:December | 0.084 | 0.016 |

Table S4: Model summary for travelling protocol birding at the national level. Numerical years refer to "migratory years". Random effect variance of Observer ID is 0.876. See model specifications described earlier.

| Fixed effect | Estimate | Standard error |
| --- | --- | --- |
| (Intercept) | -0.086 | 0.015 |
| 2019 | 0.092 | 0.016 |
| 2020 | 0.069 | 0.016 |
| 2021 | 0.029 | 0.015 |
| 2018:February | -0.071 | 0.016 |
| 2019:February | -0.052 | 0.015 |
| 2020:February | -0.166 | 0.013 |
| 2021:February | -0.066 | 0.012 |
| 2018:March | 0.040 | 0.017 |
| 2019:March | -0.524 | 0.017 |
| 2020:March | -0.081 | 0.014 |
| 2021:March | -0.029 | 0.013 |
| 2018:April | -0.018 | 0.019 |
| 2019:April | -1.645 | 0.022 |
| 2020:April | -0.252 | 0.015 |
| 2021:April | -0.026 | 0.014 |
| 2018:May | 0.040 | 0.019 |
| 2019:May | -0.928 | 0.019 |
| 2020:May | -0.644 | 0.017 |
| 2021:May | -0.050 | 0.014 |
| 2018:June | -0.036 | 0.023 |
| 2019:June | -0.086 | 0.020 |
| 2020:June | -0.472 | 0.017 |
| 2021:June | -0.407 | 0.016 |
| 2018:July | 0.028 | 0.023 |
| 2019:July | -0.156 | 0.021 |
| 2020:July | -0.468 | 0.018 |
| 2021:July | -0.219 | 0.016 |
| 2018:August | 0.019 | 0.023 |
| 2019:August | -0.087 | 0.021 |
| 2020:August | -0.414 | 0.018 |
| 2021:August | -0.178 | 0.016 |
| 2018:September | 0.095 | 0.021 |
| 2019:September | -0.054 | 0.019 |
| 2020:September | -0.260 | 0.017 |
| 2021:September | -0.119 | 0.016 |
| 2018:October | 0.105 | 0.019 |
| 2019:October | -0.008 | 0.018 |
| 2020:October | -0.157 | 0.015 |
| 2021:October | -0.004 | 0.014 |
| 2018:November | 0.121 | 0.018 |
| 2019:November | 0.044 | 0.017 |
| 2020:November | -0.094 | 0.015 |
| 2021:November | -0.001 | 0.014 |
| 2018:December | 0.062 | 0.018 |
| 2019:December | 0.016 | 0.017 |
| 2020:December | -0.054 | 0.015 |
| 2021:December | 0.083 | 0.014 |

Table S5: Model summary for list distance at the national level. Numerical years refer to "migratory years". Random effect variance of Observer ID is 0.463. See model specifications described earlier.

| Fixed effect | Estimate | Standard error |
| --- | --- | --- |
| (Intercept) | 0.841 | 0.011 |
| 2019 | -0.048 | 0.013 |
| 2020 | -0.090 | 0.012 |
| 2021 | -0.179 | 0.012 |
| 2018:February | -0.137 | 0.013 |
| 2019:February | -0.094 | 0.012 |
| 2020:February | -0.106 | 0.010 |
| 2021:February | -0.110 | 0.009 |
| 2018:March | 0.007 | 0.013 |
| 2019:March | -0.111 | 0.013 |
| 2020:March | -0.088 | 0.011 |
| 2021:March | -0.035 | 0.010 |
| 2018:April | 0.025 | 0.015 |
| 2019:April | -0.369 | 0.018 |
| 2020:April | -0.095 | 0.012 |
| 2021:April | 0.017 | 0.011 |
| 2018:May | 0.019 | 0.015 |
| 2019:May | -0.228 | 0.015 |
| 2020:May | -0.235 | 0.014 |
| 2021:May | -0.015 | 0.011 |
| 2018:June | 0.030 | 0.018 |
| 2019:June | 0.026 | 0.016 |
| 2020:June | -0.102 | 0.014 |
| 2021:June | -0.052 | 0.013 |
| 2018:July | 0.040 | 0.018 |
| 2019:July | -0.012 | 0.016 |
| 2020:July | -0.124 | 0.015 |
| 2021:July | -0.023 | 0.013 |
| 2018:August | 0.006 | 0.018 |
| 2019:August | -0.005 | 0.016 |
| 2020:August | -0.109 | 0.014 |
| 2021:August | -0.025 | 0.012 |
| 2018:September | 0.016 | 0.016 |
| 2019:September | 0.008 | 0.015 |
| 2020:September | -0.085 | 0.014 |
| 2021:September | 0.006 | 0.012 |
| 2018:October | 0.062 | 0.014 |
| 2019:October | 0.004 | 0.014 |
| 2020:October | -0.049 | 0.012 |
| 2021:October | 0.051 | 0.011 |
| 2018:November | 0.073 | 0.014 |
| 2019:November | 0.029 | 0.013 |
| 2020:November | -0.018 | 0.011 |
| 2021:November | 0.052 | 0.011 |
| 2018:December | 0.088 | 0.014 |
| 2019:December | 0.045 | 0.013 |
| 2020:December | 0.003 | 0.012 |
| 2021:December | 0.080 | 0.011 |

Table S6: Model summary for list duration at the national level. Numerical years refer to "migratory years". Random effect variance of Observer ID is 0.784. See model specifications described earlier.

| Fixed effect | Estimate | Standard error |
| --- | --- | --- |
| (Intercept) | 3.953 | 0.009 |
| 2019 | -0.020 | 0.008 |
| 2020 | -0.024 | 0.008 |
| 2021 | -0.113 | 0.008 |
| 2018:February | -0.146 | 0.008 |
| 2019:February | -0.096 | 0.008 |
| 2020:February | -0.157 | 0.007 |
| 2021:February | -0.141 | 0.006 |
| 2018:March | -0.031 | 0.009 |
| 2019:March | -0.215 | 0.008 |
| 2020:March | -0.103 | 0.007 |
| 2021:March | -0.084 | 0.007 |
| 2018:April | -0.007 | 0.009 |
| 2019:April | -0.325 | 0.008 |
| 2020:April | -0.117 | 0.008 |
| 2021:April | -0.061 | 0.007 |
| 2018:May | -0.025 | 0.010 |
| 2019:May | -0.267 | 0.008 |
| 2020:May | -0.233 | 0.008 |
| 2021:May | -0.109 | 0.007 |
| 2018:June | -0.051 | 0.011 |
| 2019:June | -0.012 | 0.010 |
| 2020:June | -0.163 | 0.008 |
| 2021:June | -0.093 | 0.008 |
| 2018:July | -0.075 | 0.012 |
| 2019:July | -0.085 | 0.010 |
| 2020:July | -0.155 | 0.009 |
| 2021:July | -0.047 | 0.008 |
| 2018:August | -0.073 | 0.011 |
| 2019:August | -0.096 | 0.010 |
| 2020:August | -0.144 | 0.008 |
| 2021:August | -0.066 | 0.008 |
| 2018:September | -0.022 | 0.010 |
| 2019:September | -0.102 | 0.010 |
| 2020:September | -0.114 | 0.009 |
| 2021:September | -0.049 | 0.008 |
| 2018:October | 0.037 | 0.009 |
| 2019:October | -0.017 | 0.009 |
| 2020:October | -0.079 | 0.008 |
| 2021:October | -0.023 | 0.007 |
| 2018:November | 0.106 | 0.009 |
| 2019:November | 0.018 | 0.009 |
| 2020:November | -0.015 | 0.008 |
| 2021:November | 0.007 | 0.007 |
| 2018:December | 0.096 | 0.009 |
| 2019:December | 0.046 | 0.009 |
| 2020:December | 0.022 | 0.008 |
| 2021:December | 0.051 | 0.007 |

Table S7: Model summary for list length at the national level. Numerical years refer to "migratory years". Random effect variance of Observer ID is 0.539. See model specifications described earlier.

| Fixed effect | Estimate | Standard error |
| --- | --- | --- |
| (Intercept) | 2.532 | 0.008 |
| 2019 | -0.004 | 0.008 |
| 2020 | 0.065 | 0.007 |
| 2021 | 0.010 | 0.007 |
| 2018:February | -0.083 | 0.007 |
| 2019:February | -0.067 | 0.007 |
| 2020:February | -0.117 | 0.006 |
| 2021:February | -0.129 | 0.006 |
| 2018:March | -0.020 | 0.008 |
| 2019:March | -0.087 | 0.007 |
| 2020:March | -0.079 | 0.007 |
| 2021:March | -0.098 | 0.006 |
| 2018:April | -0.059 | 0.009 |
| 2019:April | -0.170 | 0.007 |
| 2020:April | -0.091 | 0.007 |
| 2021:April | -0.109 | 0.007 |
| 2018:May | -0.068 | 0.009 |
| 2019:May | -0.147 | 0.007 |
| 2020:May | -0.168 | 0.007 |
| 2021:May | -0.143 | 0.007 |
| 2018:June | -0.088 | 0.010 |
| 2019:June | -0.087 | 0.009 |
| 2020:June | -0.126 | 0.008 |
| 2021:June | -0.077 | 0.007 |
| 2018:July | -0.153 | 0.011 |
| 2019:July | -0.155 | 0.010 |
| 2020:July | -0.131 | 0.008 |
| 2021:July | -0.054 | 0.007 |
| 2018:August | -0.140 | 0.010 |
| 2019:August | -0.181 | 0.010 |
| 2020:August | -0.163 | 0.008 |
| 2021:August | -0.114 | 0.007 |
| 2018:September | -0.075 | 0.009 |
| 2019:September | -0.138 | 0.009 |
| 2020:September | -0.143 | 0.008 |
| 2021:September | -0.109 | 0.007 |
| 2018:October | -0.043 | 0.009 |
| 2019:October | -0.091 | 0.008 |
| 2020:October | -0.077 | 0.007 |
| 2021:October | -0.082 | 0.007 |
| 2018:November | 0.023 | 0.009 |
| 2019:November | -0.041 | 0.008 |
| 2020:November | -0.037 | 0.007 |
| 2021:November | -0.053 | 0.007 |
| 2018:December | 0.015 | 0.008 |
| 2019:December | -0.019 | 0.008 |
| 2020:December | -0.013 | 0.007 |
| 2021:December | -0.009 | 0.007 |

Table S8: Model summary for urban birding at the national level. Numerical years refer to "migratory years". Random effect variance of Observer ID is 3.159. See model specifications described earlier.

| Fixed effect | Estimate | Standard error |
| --- | --- | --- |
| (Intercept) | -1.208 | 0.021 |
| 2019 | -0.084 | 0.020 |
| 2020 | -0.092 | 0.019 |
| 2021 | -0.023 | 0.019 |
| 2018:February | 0.069 | 0.019 |
| 2019:February | 0.129 | 0.019 |
| 2020:February | 0.293 | 0.016 |
| 2021:February | 0.077 | 0.016 |
| 2018:March | -0.024 | 0.021 |
| 2019:March | 0.445 | 0.019 |
| 2020:March | 0.099 | 0.018 |
| 2021:March | -0.035 | 0.017 |
| 2018:April | 0.052 | 0.022 |
| 2019:April | 0.996 | 0.019 |
| 2020:April | 0.225 | 0.019 |
| 2021:April | -0.054 | 0.018 |
| 2018:May | -0.227 | 0.022 |
| 2019:May | 0.755 | 0.019 |
| 2020:May | 0.635 | 0.018 |
| 2021:May | -0.080 | 0.018 |
| 2018:June | 0.052 | 0.025 |
| 2019:June | 0.073 | 0.023 |
| 2020:June | 0.572 | 0.019 |
| 2021:June | 0.486 | 0.019 |
| 2018:July | 0.168 | 0.026 |
| 2019:July | 0.232 | 0.024 |
| 2020:July | 0.515 | 0.020 |
| 2021:July | 0.223 | 0.019 |
| 2018:August | 0.127 | 0.025 |
| 2019:August | 0.158 | 0.024 |
| 2020:August | 0.447 | 0.020 |
| 2021:August | 0.114 | 0.019 |
| 2018:September | 0.154 | 0.024 |
| 2019:September | 0.185 | 0.023 |
| 2020:September | 0.412 | 0.020 |
| 2021:September | 0.042 | 0.019 |
| 2018:October | 0.061 | 0.022 |
| 2019:October | 0.040 | 0.022 |
| 2020:October | 0.291 | 0.018 |
| 2021:October | 0.033 | 0.018 |
| 2018:November | 0.014 | 0.022 |
| 2019:November | 0.047 | 0.021 |
| 2020:November | 0.220 | 0.018 |
| 2021:November | 0.047 | 0.018 |
| 2018:December | -0.127 | 0.022 |
| 2019:December | -0.144 | 0.021 |
| 2020:December | 0.103 | 0.018 |
| 2021:December | -0.085 | 0.018 |

Table S9: Model summary for spatial coverage at the national level. Numerical years refer to "migratory years". Random effect variance of Cell ID is 10.22. See model specifications described earlier.

| Fixed effect | Estimate | Standard error |
| --- | --- | --- |
| (Intercept) | -4.629 | 0.072 |
| 2019 | 0.397 | 0.073 |
| 2020 | 0.680 | 0.072 |
| 2021 | 0.768 | 0.071 |
| 2018:February | -0.058 | 0.077 |
| 2019:February | -0.079 | 0.071 |
| 2020:February | 0.035 | 0.066 |
| 2021:February | 0.435 | 0.063 |
| 2018:March | -0.257 | 0.079 |
| 2019:March | -0.376 | 0.073 |
| 2020:March | -0.252 | 0.068 |
| 2021:March | -0.090 | 0.066 |
| 2018:April | -0.568 | 0.082 |
| 2019:April | -1.190 | 0.082 |
| 2020:April | -0.610 | 0.071 |
| 2021:April | -0.423 | 0.068 |
| 2018:May | -0.620 | 0.082 |
| 2019:May | -0.934 | 0.079 |
| 2020:May | -1.088 | 0.076 |
| 2021:May | -0.484 | 0.069 |
| 2018:June | -1.344 | 0.092 |
| 2019:June | -1.275 | 0.083 |
| 2020:June | -1.232 | 0.078 |
| 2021:June | -1.222 | 0.076 |
| 2018:July | -1.457 | 0.094 |
| 2019:July | -1.465 | 0.086 |
| 2020:July | -1.542 | 0.081 |
| 2021:July | -1.099 | 0.075 |
| 2018:August | -1.481 | 0.094 |
| 2019:August | -1.614 | 0.088 |
| 2020:August | -1.361 | 0.079 |
| 2021:August | -1.017 | 0.074 |
| 2018:September | -1.111 | 0.089 |
| 2019:September | -1.210 | 0.082 |
| 2020:September | -1.243 | 0.078 |
| 2021:September | -0.861 | 0.072 |
| 2018:October | -0.656 | 0.083 |
| 2019:October | -0.799 | 0.077 |
| 2020:October | -0.716 | 0.072 |
| 2021:October | -0.414 | 0.068 |
| 2018:November | -0.408 | 0.080 |
| 2019:November | -0.513 | 0.074 |
| 2020:November | -0.344 | 0.069 |
| 2021:November | -0.405 | 0.068 |
| 2018:December | -0.228 | 0.078 |
| 2019:December | -0.322 | 0.073 |
| 2020:December | -0.335 | 0.069 |
| 2021:December | -0.188 | 0.067 |
